# Tachykinin-related peptide signalling is important for the immune response of the mealworm beetle *Tenebrio molitor* L

**DOI:** 10.1101/2025.08.21.671474

**Authors:** N. Konopińska, K. Walkowiak-Nowicka, G. Nowicki, M. Keshavarz, Sz. Chowański, J. Rolff, A. Urbański

**Author notes:** Corresponding author: Natalia Konopińska.

## Abstract

Insects possess a well-developed innate immune system, which encompasses both cellular and humoral mechanisms. On the basis of the similarities in neuropeptide actions between insects and vertebrates, we assume that neuropeptides such as tachykinin-related peptides (TRPs) regulate insect immune responses and are themselves modulated following infection. In this study, we examined how immune activation affects the expression of genes encoding TRP precursors and receptors (*TRP* and *TRPR*) and whether TRPs directly modulates selected immune mechanisms in the pest species *Tenebrio molitor*.

Our results revealed two important insights. First, after activation of the immune system, *TRP* and *TRPR* genes were significantly downregulated in the nervous system and immune-related cells. These changes are closely correlated with the changes of the expression level of immune genes. We then show using Spantide II, a potent antagonist of TRPR, and RNAi knock-down of *TRP* and *TRPR* the modulation of key processes of the *T. molitor* humoral response. This includes the over-expression of genes encoding antimicrobial peptides and the important arthropod immune effector phenoloxidase activity.

Our findings highlight a compelling association between the TRP and immune regulation in *Tenebrio* and provide insights into the hormonal regulation of physiological processes in insects. Our research also provides novel insights that can contribute to the development of sustainable pest control strategies amid increasing insecticide resistance.

## 1. Introduction

Insects are the most diverse taxon on earth, and they have a significant impact on our lives. Their roles in pollination and sanitation, for example, are crucial to human well-being. But insects as pests also cause substantial economic losses [1]. This situation necessitates the development of new, biosafe, and targeted agents for pest control. Paradoxically, the mass rearing of certain pest species, such as *Tenebrio molitor*, can help meet the growing food demands of the human population [2]. For this reason, a better understanding of the immunology of *T. molitor* is essential not only for developing new pest control strategies but also for improving insect mass-rearing practices.

Insects have an innate immune system, which is composed of cellular and humoral responses, resulting in effective protection against infection [3]. The cellular response entails processes such as phagocytosis, nodulation, and encapsulation, in which haemocytes - the main cells of the insect haemolymph - are involved. The humoral response comprises the activity of proteins and enzymes such as lysozyme, antimicrobial peptides (AMPs), and phenoloxidase (PO), an important enzyme in arthropod immune function that generates cytotoxic substances but also contributes to the sclerotization of the cuticle [4]. However, the activation and coordination of these responses require precise regulation. Emerging evidence suggests that this regulation is at least partially mediated by neuroendocrine factors, including neuropeptides that can directly interact with immune cells [3,5]. Neuropeptides are key regulators of physiological homeostasis and are involved in all aspects of insect biology, including development, metabolism, behaviour, and reproduction. Importantly, recent studies indicate that they may also act as immunomodulators, highlighting their role in the cross-talk between the neuroendocrine and immune systems. These small signalling molecules are primarily produced and secreted by neurosecretory cells (NSCs) in the insect central nervous system, but endocrine cells in peripheral tissues such as the gut and reproductive system also contribute to their production [6].

One of the largest and most important families of neuropeptides identified in insects is the tachykinin-related peptides (TRPs). They show high structural and functional homology with tachykinins (TKs) present in vertebrates [7]. Tachykinins such as vertebrate substance P (SP) have pleiotropic effects and take part in nociception, stress responses, and the regulation of muscle contractions [7]. These functions are consistent with the action of TRPs in insects [7,8]. Moreover, both neuropeptide families interact with specific G protein-coupled receptors (GPCRs), leading to the activation of signalling pathways affecting cell activity [9]. Our previous studies indicated that TRPs can influence both the cellular response by modulating haemocyte morphology and activity and the humoral response by regulating the expression levels of immune-related genes in insects [4,10]. This finding is in line with the effect of SP on the activity of the vertebrate immune system [11]. Moreover, our previous studies showed that human SP influences the immune system activity of *Tenebrio molitor* beetles, which supports the hypothesis that the immunomodulatory role of the TK system is evolutionarily conserved [10]. Given the key role that neuropeptides play in signalling within the nervous system and other tissues, it seems likely that TRPs influence the process of pathogen recognition and modulate the overall immune response. The mechanisms, however, by which TRPs regulate the immune response remain largely unknown. Therefore, here, we study two essential questions to investigate the immunoregulatory role of TRPs in insects. First, whether the TRP pathway is modulated during the immune response against bacterial components or after cytokines administration? Second, are TRPs directly involved in the regulation of the insect immune system?

The first part of our research focused on evaluating changes in the expression patterns of genes encoding TRP precursor (*TRP*) and receptor (*TRPR*) in different phases of immune system activation. To obtain a more complete picture of the changes observed in the TRP system, the *T. molitor* immune system was activated *via* different activators, including one of the most important insect cytokines, Spätzle-like (*Tm*Spz-like). The use of *Tm*Spz-like will allow us to better support our previous finding that cytokines are modulators of the neuroendocrine system in insects, similar to humans [12]. Because of the nature of neuropeptides, which are most likely stored in neuroendocrine cells and released during a specific physiological state, molecular analysis of their expression levels was supported by immunocytochemical analysis of the presence of TRP precursors in the *Tenebrio* nervous system. This new finding strongly suggests that insect cytokines can affect neuropeptide levels in insect neuroendocrine cells.

In the second part of the research, we focused on the direct effect of TRP on the *T. molitor* immune system on the basis of the expression of immune-related genes and phenoloxidase. For this purpose, we used RNAi gene knock-down and Spantide II, a potent antagonist of TRPR. Using both dsRNA to selectively disrupt TRP-related pathways and Spantide II to block TRPRs will provide a comprehensive understanding of TRP function and its impact on immune system modulation. Additionally, we identified the combination that most significantly affect *T. molitor* immune system function and then tested the survival of these groups following an additional application of *Escherichia coli*, which, under normal conditions, significantly reduces beetle survival.

Our research advances the understanding of how the TRP system influences immune regulation in insects. These findings may support the development of more effective pest control strategies and improve insect mass rearing methods. By uncovering how hormonal pathways affect immunity, we identify potential targets for pest management and ways to enhance the health and productivity of farmed insects used in food and feed.

## 2. Results

### 2.1. Expression patterns of the *TRP* and *TRPR* genes after activation of the *Tenebrio* immune system

To evaluate the potential participation of TRP signalling in the immune response after *T. molitor* immune system activation, the expression levels of genes encoding TRP precursors (*TRP*) and the receptor (*TRPR*) were analysed in the nervous system (brain and ventral nerve cord (VNC)) and immune-related cells (fat body and haemocytes). The analysis of three time points (3, 6, and 24 hours) revealed changes in *TRP* and *TRPR* gene expression at different stages of the immune response [12,13]. To activate different signalling pathways involved in immune system activation, insects were injected with lyophilized *Escherichia coli* (*Ec)*, and peptidoglycan of *Staphylococcus aureus* (PG). Additionally, to evaluate the potential role of insect cytokines in the modulation of the insect immune system, beetles were also injected with *Tm*Spz-like, a key cytokine and ligand of the Toll receptors in insects [14]. The molecular analysis was preceded by survival experiments, which revealed significant differences between beetles injected with different activators of the immune system (Fig. 1A). After *Ec* treatment, the survival of *T. molitor* males was significantly lower than that of control individuals injected with physiological saline (PS) (Gehan–Breslow–Wilcoxon test, *p* ≤ 0.01, n=50 per treatment). Moreover, statistically significant differences between survival curves were reported between *Tm*Spz-like treatment and *Ec* (Gehan-Breslow-Wilcoxon test, *p* ≤ 0.05, n=50 per treatment).

**Fig. 1.**
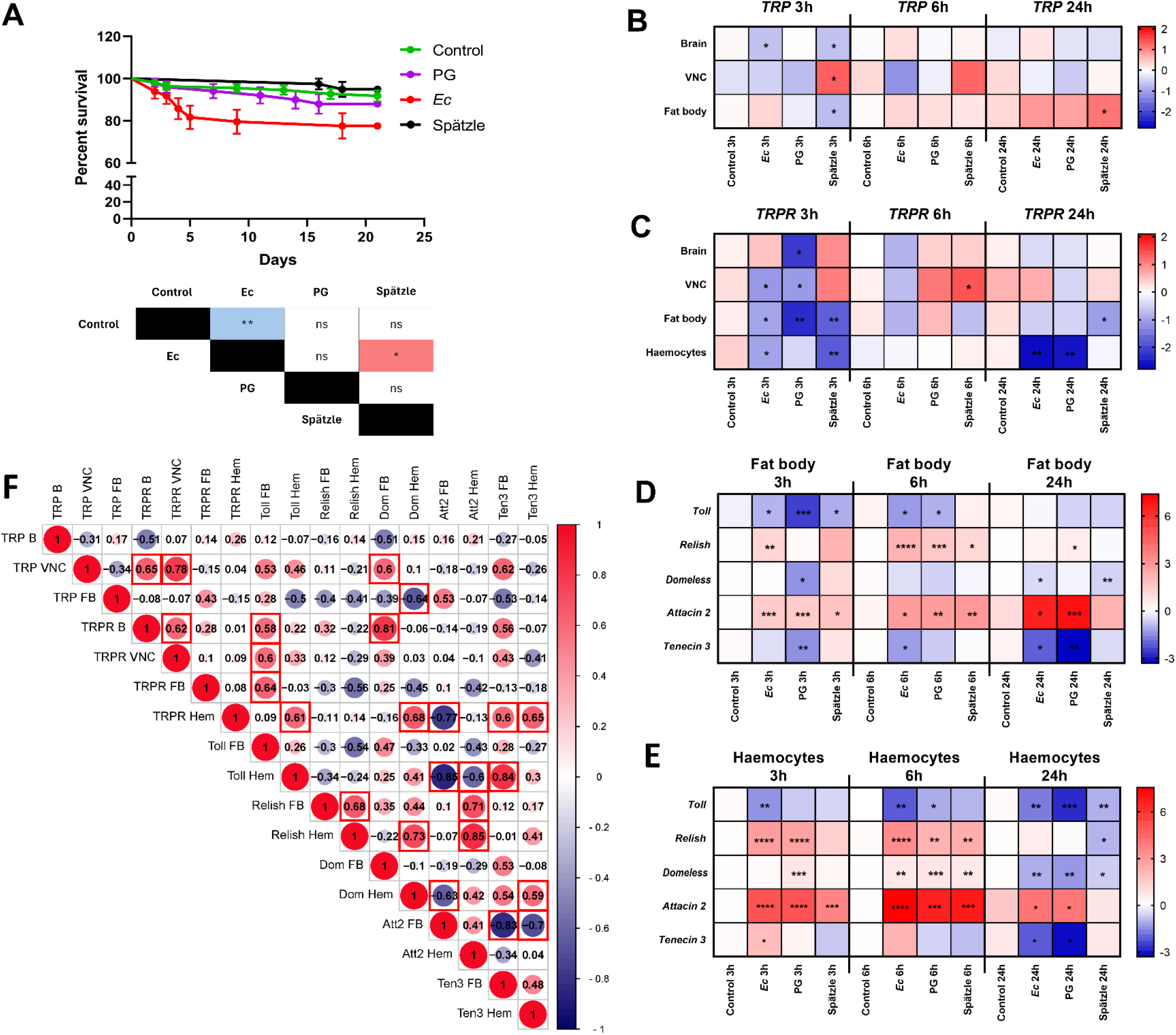
Expression patterns of the TRP and TRPR genes after activation of the *Tenebrio* immune system according to additional analyses. **A**. Survival curve of *T. molitor* males after injection of physiological saline (control, green line) or peptidoglycan from *Staphylococcus aureus* (PG, 1 mg/mL, OD_600_ = 1, violet line), a suspension of lyophilized *Escherichia coli* K12 (*Ec*, 1 mg/mL, OD_600_ = 1; red line), or a suspension of *Tm*Spz-like at a concentration of 10^−7^ M (black line). The colours in the table indicate an increase (red) or decrease (blue) in survival relative to the group in the top row of the table. The values are presented as the means ± SEs. Table – Statistical comparison of estimated survival curves based on the Gehan-Breslow‒ Wilcoxon test; ns – nonsignificant difference, \**p* ≤ 0.05, \*\**p* ≤ 0.01, n = 50 per research variant. **B-E**. Heatmaps showing the changes in the expression levels of genes encoding TRP precursor (B) and TRP receptor (TRPR) (C) in the brain, ventral nerve cord (VNC), fat body, and haemocytes and the changes in the expression levels of immune-related genes in the fat body (D) and haemocytes (E) 3, 6 and 24 hours after the application of physiological saline (control), *Ec*, PG or *Tm*Spz-like protein. The values are expressed as log2fold values, shades of red indicate upregulation, shades of blue indicate downregulation, \**p* ≤0.05, \*\**p* ≤0.01, *** *p* ≤0.001, **** *p* ≤0.0001, n = 3 per research variant, one biological repetition was samples collected from at least 20 (B, VNC, haemocytes) or 10 (fat body) individuals. **F -** Correlation matrix showing the dependencies between the expression levels of TRP and TRPR genes in the brain (B), ventral nerve cord (VNC), fat body (FB), and haemocytes (Hem). These changes were correlated with the expression levels of immune-related genes (*Toll, Relish, Domeless* (Dom), *Attacin 2* (Att2), and *Tenecin 3* (Ten3)) in the fat body (FB) and haemocytes (Hem). To estimate the correlation of the data, the Pearson correlation coefficient method was used. The matrix was generated via SRplot software (https://www.bioinformatics.com.cn/srplot). Size of the dot - level of the r value. The r value is presented in the middle of the dot. Different colours indicate different r values. Red shading indicates positive correlations (r ≥ 0), and blue shading indicates negative correlations (r ≤ 0). Red squares indicate statistically significant correlations (*p* ≤ 0.05).

The analysis of the expression patterns of *TRP* revealed that the significant changes in expression level were mostly visible at the first tested time point (Fig. 1 and Fig. S1). After 3 h, a significant decrease in the expression level of the *TRP* gene in the fat body and brain of beetles injected with *Tm*Spz-like protein was observed (Fig. 1B and Fig. S1). Significant downregulation of *TRP* was also observed in the brain after *Ec* application. Significant overexpression of *TRP* was only visible in the VNC after *Tm*Spz-like protein application. At later time point, 6 h after the application of immune system activators, no significant changes were observed. After 24 h, the only significant increase that occurred was in the fat body after *Tm*Spz-like protein application.

In the case of *TRPR*, a greater number of significant changes were observed (Fig. 1C and Fig. S2). After 3 h, a decrease in the expression level of the *TRPR* gene was detected in the brain (after the application of PG), VNC (*Ec* and PG), fat body (all groups), and haemocytes (*Ec* and *Tm*Spz-like protein). The only increase in the expression level was detected in VNC 6 hours after the application of the *Tm*Spz-like protein. However, after 24 hours, there was a decrease in the expression level of *TRPR* in the fat body (after *Tm*Spz-like protein) and haemocytes (after *Ec* and PG).

Next we examined changes in the expression levels of selected immune-related genes in cells directly involved in the immune response (fat body cells and haemocytes) and correlated with changes in the expression levels of the *TRP* and *TRPR* genes after immune activation (Fig. 1D, 1E and Fig. S3, S4). In the fat body, in the 3-hour variant, a significant decrease in the expression level of genes encoding Toll was detected in all the tested groups. The expression of *Domeless* and *Tenecin 3* genes also decreased 3 h after PG application. A significant increase in expression level was detected for *Relish* (*Ec*) and *Attacin 2* (all groups). At six hours, there was a significant increase in the expression levels of *Relish* and *Attacin 2* and a decrease in the expression levels of *Toll* (*Ec* and PG) and *Tenecin 3* (*Ec*). Twenty-four hours after the application of the immune system activators, *Relish* (PG) and *Attacin 2* (*Ec* and PG) were overexpressed. A decrease in the expression level occurred in *Domeless* after the application of *Ec* and *Tm*Spz-like protein. There was also a decrease in the expression level of genes encoding *Tenecin 3* after *Ec* and PG injection at that time.

In haemocytes, 3 hours after immune activation, there was a significant decrease in the expression level of the *Toll* gene (after *Ec* application). Moreover, significant overexpression of the genes encoding *Domeless* (PG), *Relish* (*Ec*, PG), *Attacin 2* (all groups) and *Tenecin 3* (*Ec*) was detected (Fig. 3). After 6 h, there was a significant increase in the expression level of genes encoding *Relish, Domeless* and *Attacin 2* in all the tested groups. On the other hand, *Toll* (*Ec* and PG) was significantly downregulated. After 24 hours, only an increase in the expression level of *Attacin 2* (*Ec* and PG) was observed. Additionally, in the case of haemocytes, we observed the downregulation of *Toll* and *Domeless* after the application of each type of activator. In contrast, the gene encoding *Relish* was significantly downregulated only after *Tm*Spz-like protein injection. Moreover, a decrease in the expression level of the gene encoding *Tenecin 3* was detected after *Ec* and PG application.

We found a significant positive correlation between the expression of *TRP* in VNC and the *TRPR* in the brain and VNC. The same correlation also occurred in *TRPR* between the brain and VNC (Fig. 1F). Interestingly, some dependencies were detected between the expression levels of genes encoding *TRP, TRPR* and immune-related genes (Fig. 1F). A significant positive correlation has been reported between the expression of *Toll* in the fat body and *TRPR* in the brain, VNC, and fat body. Moreover, the expression level of *Domeless* was positively correlated with the expression of *TRP* in the VNC and *TRPR* in the brain. Conversely, the expression of *Domeless* in haemocytes is negatively correlated with *TRP* expression in the fat body. When changes in the level of *TRPR* expression in haemocytes were analysed, a positive correlation was observed between *TRPR* and *Toll, Domeless* and *Tenecin 3*. On the other hand, in haemocytes, a negative correlation between the expression level of *TRPR* and that of *Attacin 2* was also detected.

Significant correlations were also reported between the expression of genes related to immune system activity (Fig. 1F). Interestingly, the expression of *Toll* in the fat body was not correlated with other immune genes. In contrast, *Toll* expression in haemocytes was significantly negatively correlated with *Attacin 2* expression in both the fat body and haemocytes. However, the expression of *Toll* was positively correlated with *Tenecin 3* expression in the fat body. Some correlations between the expression levels of *Relish* in fat body and haemocytes have also been reported. In the fat body, the *Relish* expression level was positively correlated with the expression of genes encoding Relish and Attacin 2 in haemocytes. However, strong positive correlations between the expression of *Relish* and that of *Domeless* and *Attacin 2* in haemocytes were observed. Moreover, in haemocytes, the expression level of *Domeless* was negatively correlated with the expression of *Attacin 2* and positively correlated with *Tenecin 3*. In the case of direct correlations between genes encoding the tested AMPs, the expression of fat body *Attacin 2* was negatively correlated with the expression level of *Tenecin 3* in the fat body and haemocytes.

### 2.2. Immunolocalization of TRP precursor in the nervous system after activation of the immune system

To analyse changes in the distribution of the TRP precursor following immune system activation, immunolocalization was performed. The immunocytochemical results revealed differences in the distribution and abundance of the TRP precursor in both the brain and different ganglions of the VNC of *T. molitor*. These changes were time- and immune activator dependent. After 3 h (Fig. 2A), a relative decrease in the intensity and abundance of the fluorescent signal associated with the immunolocalization of TRP was observed in all the parts of the VNC after *Ec* injection. A weaker fluorescence signal also occurred in the second abdominal ganglion after the application of PG and the *Tm*Spz-like protein and in the terminal abdominal ganglion (TAG) after the application of the *Tm*Spz-like protein. After 6 h in the group activated with *Ec*, a decrease was observed in all the examined tissues, including the brain (Fig. 2B). After PG application, the signal intensity and abundance were relatively greater than those in the control group. Additionally, in the brain after the injection of *Tm*Spz-like protein, such an increase occurred with a simultaneous decrease in the relative amount of the precursor in the second abdominal ganglion and TAG. Long-term effects (24 h incubation) were observed only after *Ec* application in the second terminal ganglion and the second abdominal ganglion (Fig. 2C).

**Fig. 2.**
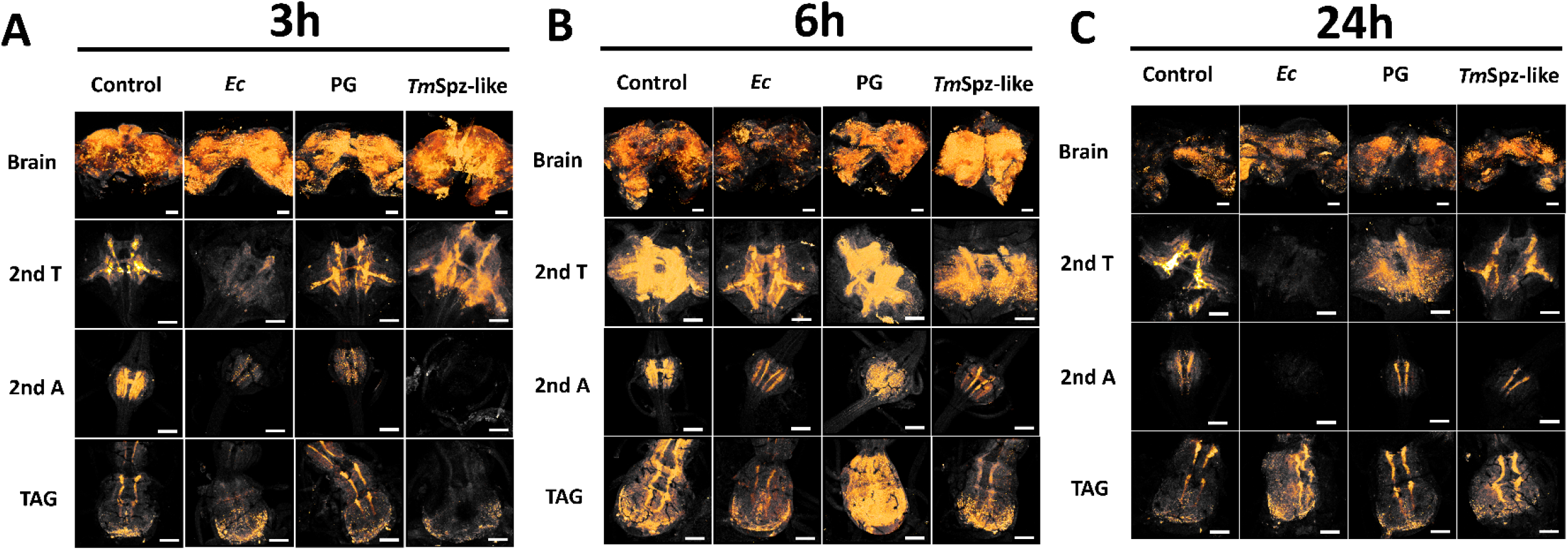
Micrographs showing the distribution of TRP precursor in the brain; second terminal ganglion (2^nd^ T), second abdominal ganglion (2^nd^ A) and terminal abdominal complex (TAG) 3 (A), 6 (B) and 24 h (C) after the application of physiological solution (control); *Escherichia coli* (*Ec*), *Staphylococcus aureus* peptidoglycan (PG) and Spätzle-like protein (*Tm*Spz-like). The orange colour shows a 3D projection of the presented structure, which was superimposed on the pictures to show the distribution of the TRP precursor. A 3D projection was created with AMIRA 3D software on the basis of the obtained Z-stack files of the whole analysed structure (threshold 75-255 for fluorescent intensity). Sample examination was performed on the same day under the same conditions and microscope setup, which allowed comparison of the obtained micrographs. Scale (white line) 100 µm.

**Fig. 3.**
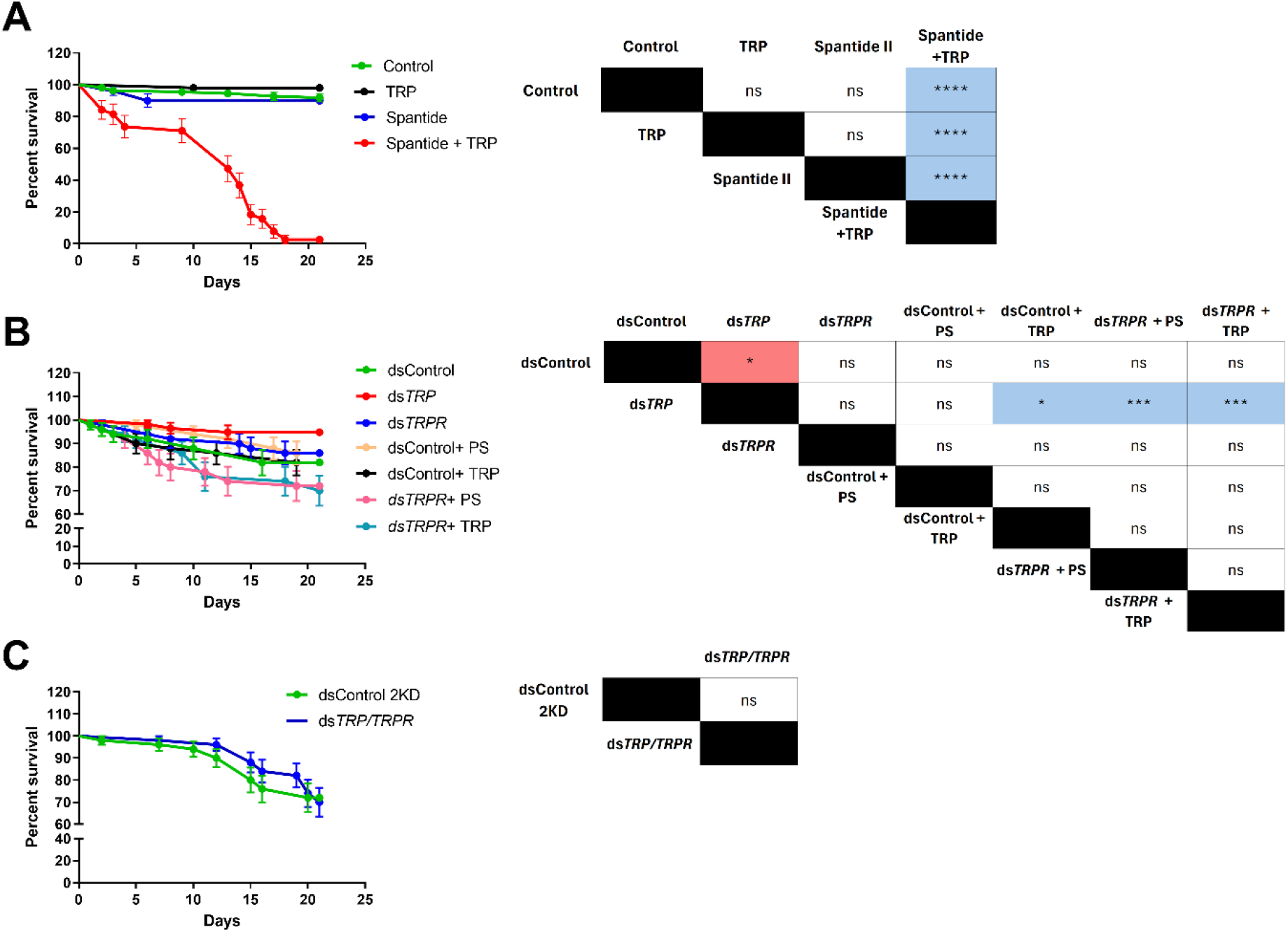
Survival curves of *T. molitor* males after injection of a potent antagonist of the TRP receptor Spantide II (A), knockdown of the *TRP* and *TRPR* genes (B), and double knockdown of the *TRP* and *TRPR* genes (C). In graph A, control represents individuals injected with physiological saline; TRP represents individuals injected with Tenmo-TRP-7 at a concentration of 10^-5^ M; Spantide II represents beetles injected with Spantide II at a concentration of 10^-3^ M; and Spantide + TRP represents simultaneous injection of Spantide II and Tenmo-TRP-7 at a concentration of 10^-5^ M. In graph B, Control individuals were injected with dsRNA targeted to *Galleria mellonella lysozyme* (*GmLys*), while gene knockdowns were performed using ds*TRP* for the *TRP* gene and ds*TRPR* for the TRP receptor gene (*TRPR*). The dsControl + PS represents insects injected with *dsGmLys* and physiological saline; dsControl + TRP represents individuals injected with *dsGmLys* and Tenmo-TRP-7 at a concentration of 10^-5^ M; dsTRPR + PS represents insects injected with *dsTRPR* and physiological saline; dsTRPR + TRP represents insects injected with *dsTRPR* and Tenmo-TRP-7 at a concentration of 10^-5^ M. In Graph C, Control 2KD individuals were injected with a double dose of ds*LysGm*, while ds*TRP/TRPR* was used for the double knockdown of *TRP* and *TRPR*. The colours in the table indicate an increase (red) or decrease (blue) in survival relative to the group in the top row of the table. The values are the means ± SEMs. Table – Statistical comparison of estimated survival curves based on the Gehan-Breslow‒Wilcoxon test; ns – nonsignificant differences, * *p*≤0.05, ** *p*≤0.01, *** *p*≤0.001, **** *p*≤0.0001, n = 50 per research variant.

### 2.3. Evaluation of the potential direct effects of TRP on insect immune mechanisms

In previous experiments, we reported that the expression of immune-related genes in *T. molitor* was associated with changes in the expression levels of *TRP* and *TRPR*. However, it remains unclear whether the observed immune responses are directly dependent on TRP signalling and thus whether TRPs play a direct role in regulating immune function in insects. Therefore, we used Spantide II to pharmacologically inhibit TRPR activity. We also applied dsRNA to knockdown the genes encoding the TRP precursor and receptor (consequently, ds*TRP* and ds*TRPR*). This combined pharmacological and molecular approach enabled us to assess the direct involvement of TRP signalling in modulating immune gene expression and insect survival. Moreover, to confirm the specificity and effectiveness of TRP pathway inhibition, we readministered synthetic TRP (Tenmo-TRP-7, amino acid sequence: MPRQSGFFGMRa which possesses a sequence similar to the native TRP identified in *T. molitor*) to selected groups.

#### 2.3.1. Survival experiment after Spantide II and dsRNA treatment

First we focused on the effects of the tested compounds on the survival of *T. molitor* males. Physiological saline, Tenmo-TRP-7, and Spantide II did not affect the survival of *T. molitor* males (Fig. 3A). In these cases, around 90% of individuals survive. Simultaneous injection of Tenmo-TRP-7 and Spantide II significantly influences beetle mortality. Until the last day of observation (21^st^ day), only 2.63% of the tested individuals survived. The comparison of survival curves revealed significant differences between beetles simultaneously injected with Tenmo-TRP-7 and Spantide II and the remaining variants (Gehan-Breslow-Wilcoxon test, *p* ≤ 0.0001, in all cases, n=50 per treatment).

The survival experiment after dsRNA treatment revealed significant differences (Gehan-Breslow-Wilcoxon test, *p* ≤ 0.05, n=50 per research variant) (Fig. 3B). After *TRP* knockdown, a significant reduction in *T. molitor* mortality compared with that in the control group was observed (Gehan-Breslow-Wilcoxon test, *p* ≤ 0.05, n=50 per research variant). Interestingly, a comparison of survival curves after knockdown of the *TRP* gene revealed significant differences compared with the groups treated with ds*GmLys* (dsControl) and Tenmo-TRP-7. Significant differences compared to ds*TRP* were also observed with groups injected with ds*TRPR* and physiological saline, and ds*TRPR* and Tenmo-TRP-7 (Gehan-Breslow-Wilcoxon test, *p* ≤ 0.05 and *p* ≤ 0.001, respectively, n=50 per treatment). Compared with the control, double knockdown (ds*TRP/TRPR*) did not result in any significant differences (Fig. 3C).

#### 2.3.2. Changes in the expression levels of immune-related genes

Our previous transcriptomic analysis revealed that the application of Tenmo-TRP-7 led to significant changes in the expression patterns of a wide spectrum of immune-related genes [15]. For this reason, the next step in our research was the determination of the effects of Spantide II, ds*TRP* and ds*TRPR* on the expression levels of selected immune genes in the fat body and haemocytes. For analysis of basic immune pathways involved in the stress response, *Toll* (receptor of the Toll signalling pathway), *Relish* (transcription factor of the Imd signalling pathway), and *Domeless* (receptor of the JAK/STAT signalling pathway) were chosen. Moreover, the expression of genes encoding selected AMPs (*Attacin 2* and *Tenecin 3*) was analysed. AMP gene selection was based on transcriptomic changes occurring after Tenmo-TRP-7 application [15].

RT‒qPCR analyses revealed that changes in the expression levels of immune-related genes after the application of Spantide II and dsRNA differed depending on the tested immune tissues (Fig. 4 and Fig. S5). In the fat body, the application of Tenmo-TRP-7 decreases the expression of *Toll, Domeless* and *Attacin 2*. Injection of Spantide II and blocking of TRPR elicited effects opposite to those of Tenmo-TRP-7 injection, and *Toll* overexpression was observed. Nevertheless, Spantide II injection led to downregulation of the *Attacin 2* gene, and upregulation of *Tenecin 3* gene. The application mixture of Tenmo-TRP-7 and Spantide II abolished the effect of separate injection of Tenmo-TRP-7 and Spantide II. Interestingly, this mixture led to a decrease only in the *Attacin 2* expression level. Knockdown of the precursor and the receptor individually did not induce any changes; only the double knockdown led to a marked decrease in the expression levels of *Toll* and *Relish*. Insects treated with dsGmLys (dsControl) followed by Tenmo-TRP-7 injection showed that non-specific dsRNA did not strongly affect the action of Tenmo-TRP-7, and a significant reduction in the expression levels of *Toll, Relish, Domeless*, and *Tenecin 3* was observed. Interestingly, after receptor silencing and Tenmo-TRP-7 application, *Domeless* and *Attacin 2* were overexpressed.

**Fig. 4.**
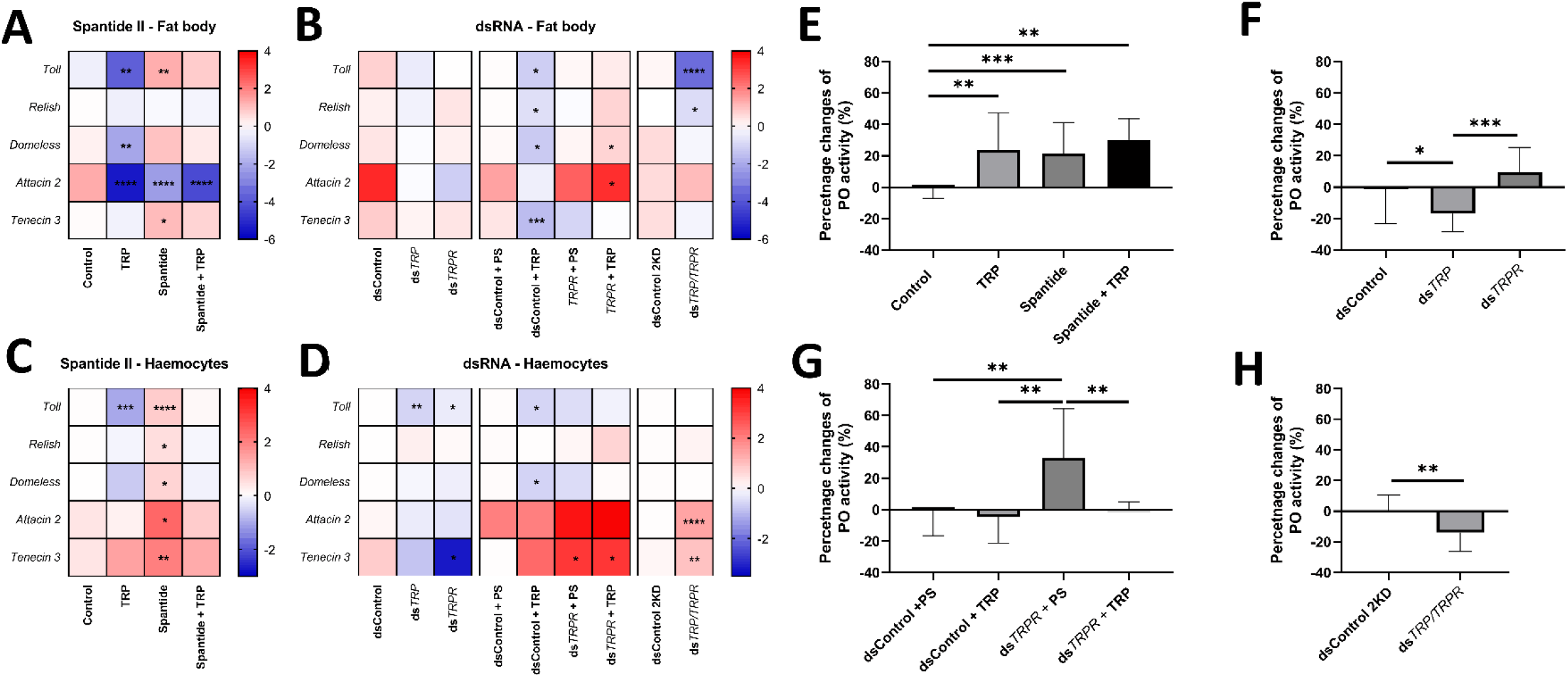
Changes in the humoral response of *T. molitor* after the application of the TRPR agonist Spantide II and the knockdown of the *TRP* and *TRPR* genes. **A-D**. Heatmaps showing changes in the expression levels of immune-related genes after Spantide II (A and C) or dsRNA treatment (B and D) in the fat body and haemocytes, respectively. Expression levels of genes tested in the fat body and haemocytes after the application of physiological saline (control), Tenmo-TRP-7 (TRP) at a concentration of 10^−5^ M, Spantide II at a concentration of 10^-3^ M, or a mixture of Tenmo-TRP-7 and Spantide II (A and C). Expression levels of immune-related genes after the injection of dsRNA-targeted genes encoding lysozyme in *Galleria mellonella* (*GmLys*, dsControl) or genes related to TRP signalling (genes encoding the TRP precursor (ds*TRP*) or receptor (ds*TRPR*)) (B and D). The effects of double injection of dsRNA and physiological saline or Tenmo-TRP-7 (TRP) at a concentration of 10^−5^ M were also tested. In addition, the effects of both TRP and TRPR knockdown were analysed. The control for the double-knockdown experiment (dsControl 2KD) was the injection of a double dose of dsRNA-targeted *GmLys*. The values are expressed as log2fold values, shades of red indicate upregulation, shades of blue indicate downregulation, \**p*≤0.05, \*\**p*≤0.01, *** *p*≤0.001, \*\*\*\**p*≤0.0001, n = 3 per research variants, as one biological repetition is considered the samples collected from 10 (fat body) or 20 (haemocytes) individuals. **E-H**. Phenoloxidase (PO) activity in *T. molitor* haemolymph after Spantide II and a mixture of Tenmo-TRP-7 treatment (**E**), knockdown of TRP and TRPR (**F**), double injection of dsRNA and physiological saline or TRP (**G**) or double knockdown of the tested genes (**H**). **A** – Phenoloxidase activity in the haemolymph of *T. molitor* after the application of physiological saline (Control), Tenmo-TRP-7 (TRP) at a concentration of 10^−5^ M, Spantide II at a concentration of 10^−3^ M, or a mixture of Tenmo-TRP-7 and Spantide II. **F**– PO activity after injection of dsRNA-targeted *Galleria mellonella* lysozyme (*GmLys*) (dsControl) or dsRNA-targeted genes related to TRP signalling (genes for TRP precursor (ds*TRP*) or receptor (ds*TRPR*)). **G** – PO activity after application of dsRNA and additional injection with physiological saline or Tenmo-TRP-7 (TRP) at a concentration of 10^−5^ M. **H** – PO activity determined after injection of a double dose of dsGmLys (dsControl 2KD) or double knockdown of *TRP* and *TRPR*. The values are presented as the percentage change relative to the control group. The results are presented as the means ± SDs, \**p*≤0.05, \*\**p*≤0.01, \*\*\**p*≤0.001, n ≥ 10 per research variant.

In haemocytes, Tenmo-TRP-7 injection significantly reduced the expression level of Toll, whereas TRPR blocking through treatment with Spantide II elicited opposite effects, with a significant upregulation of all the tested genes observed. (Fig. 4 and Fig. S6). A mixture of Spantide II and Tenmo-TRP-7 abolished the effects observed with separate injections of these compounds and did not cause any changes in the expression levels of the selected immune-related genes.. The use of dsRNA targeted to *TRP* causes a significant decrease in the expression of *Toll* in haemocytes. Knockdown of *TRPR* significantly decreases the expression levels of *Toll* and *Tenecin 3*. In contrast, simultaneous silencing of both *TRP* and *TRPR* resulted in significant overexpression of *Attacin 2* and *Tenecin 3*. Similar to a single injection of Tenmo-TRP-7, the application of ds*GmLys* and Tenmo-TRP-7 decreased the levels of *Toll* and *Domeless*, whereas in insects injected with ds*TRPR*, the administration of either physiological saline or Tenmo-TRP-7 led to *Tenecin 3* overexpression. Increased expression of *Attacin 2* and *Tenecin 3* was also detected when both the *TRP* and *TRPR* genes were silenced simultaneously.

#### 2.3.3. Phenoloxidase activity (PO)

Our previous studies revealed that Tenmo-TRP-7 significantly affects PO system activity. This was expressed as changes in PO activity and changes in the expression levels of genes related to the PO system [4]. For this reason, the next step was to analyse changes in the activity of this enzyme after Spantide II and dsRNA application. Like in previous studies, Tenmo-TRP-7 application significantly increased PO activity (Dunnett’s *post hoc* test, *p* ≤ 0.01, n≥ 10 per research variant). Blocking the receptor with Spantide II and using a mixture of Spantide II and Tenmo-TRP-7 also led to increased PO activity (Dunnett’s *post hoc* test, *p* ≤ 0.001 and *p* ≤ 0.01, respectively, n≥ 10 per research variant) (Fig. 4E).

Compared with the control, the use of dsRNA directed against the TRP precursor and simultaneous knockdown of *TRP* and *TRPR* resulted in a significant decrease in PO activity (Dunnett’s *post hoc* test, *p* ≤ 0.05; Student’s *t* test, *t*=2.90, *p* ≤ 0.01, respectively, n≥ 10 per research variant) (Fig. 4F and H). However, significant differences were also observed in PO activity between the groups in which *TRP* and *TRPR* were knocked down (Dunnett’s *post hoc* test, *p* ≤ 0.001, n≥ 10 per research variant). Interestingly, compared with the other treatments, the injection of physiological saline after ds*TRPR* treatment caused a significant increase in PO activity (Dunnett’s *post hoc* test, *p* ≤ 0.001, all cases, n≥ 10 per research variant) (Fig. 4G).

### 2.4. Survival assessment following TRP system suppression and immune activation with *Escherichia coli*

On the basis of previous analyses, we selected the experimental variants that had the strongest impact on the immune system activity of *T. molitor* to assess how the modulation of TRP signalling affects insect survival during immune system activation. For this purpose, we conducted a survival experiment in which *Ec* injection, which significantly affected *Tenebrio* survival (see above). The results revealed statistically significant changes in survival only in the group of insects with double knockdown of *TRP* and *TRPR* genes. In this group, survival was significantly greater than that in the groups injected with physiological saline, Tenmo-TRP-7, or Spantide II (Gehan-Breslow‒Wilcoxon test, *p* ≤ 0.05, in all cases, n=50 per research variants) (Fig. 5). Also during the first days of the experiment, a significant decrease in the survival of *T. molitor* treated with *Ec* and Tenmo-TRP-7 or Spantide II compared with that of the control beetles was observed (Fig. S8).

**Fig. 5.**
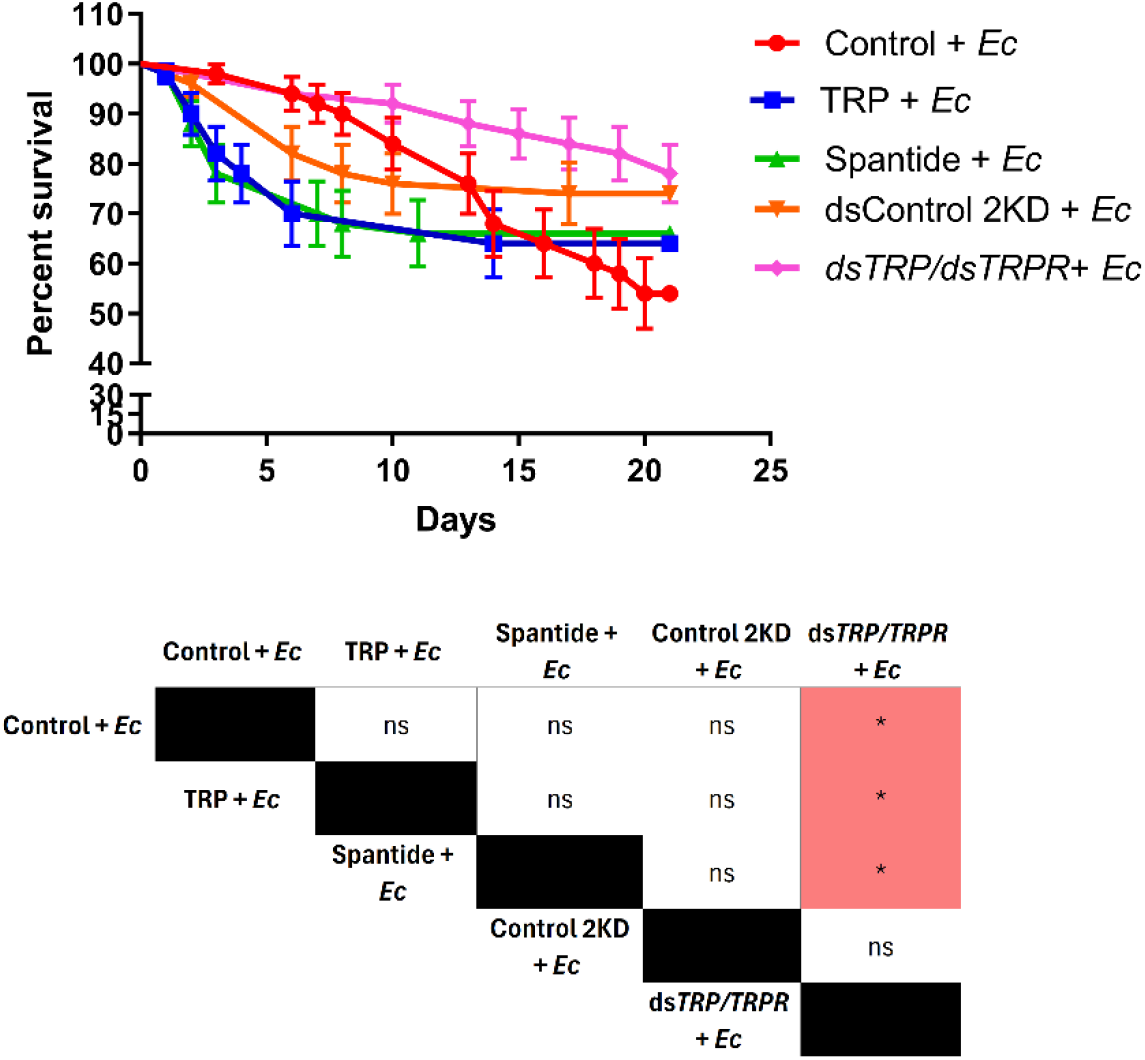
Survival curves of *T. molitor* males after injection of Tenmo-TRP-7 (TRP), the potent TRP receptor antagonist Spantide II, and double knockdown (ds*TRP/TRPR*), followed by immune system activation via *Escherichia coli* (*Ec*). Control represents individuals injected with physiological saline; TRP represents individuals injected with Tenmo-TRP-7 at a concentration of 10^− 5^ M; Spantide II represents beetles injected with Spantide II at a concentration of 10^−3^ M. After 2 hours, the groups were injected with *Ec*. For the double-knockdown control (Control 2KD) individuals were injected with a double dose of ds*LysGm*, while ds*TRP/TRPR* was used for the double knockdown of *TRP* and *TRPR*. After seven days, individuals were injected with *Ec*. The colours in the table indicate an increase (red) in survival relative to the group in the top row of the table. The values are the means ± SEMs. Table – Statistical comparison of estimated survival curves based on the Gehan-Breslow‒Wilcoxon test; ns – nonsignificant difference, * *p*≤0.05, n = 50 per research variant.

## 3. Discussion

Although our previous studies explored the influence of TRPs on immune responses, it remains unclear whether these effects are direct or indirect, highlighting the need to clarify how TRP signalling interacts with immune activation. Therefore, we first examined how the expression of genes encoding the TRP precursor and receptor changed after activation of the immune system. In humans, SP levels fluctuate depending on the physiological state of the body [16]. Given the structural and functional similarities between SP and TRP, we hypothesized that analogous changes may occur in insects. What is important, overall, the main changes were significant downregulation of *TRPR* gene in the nervous system and immune-related cells. The downregulation of *TRP* was also noted, but to a lesser extent. These findings suggest that once the immune response is triggered, TRP-related signalling can be inhibited. This notion may be associated with our previous studies looking at the immunomodulatory effect of TRP in *T. molitor*. Research by Urbański et al. [4,15] revealed that the application of Tenmo-TRP-7 significantly downregulated some immune-related genes and decreased the antimicrobial activity of *T. molitor* haemolymph. Considering these results, we may assume that TRP signalling is inhibited at the initial stage of immune system activation, probably to elicit maximal activation of immune mechanisms. Moreover, *TRP* and *TRPR* expression are significantly correlated across different parts of the nervous system. This positive correlation in the nervous system may be due to regulatory mechanisms aimed at maintaining homeostasis and ensuring proper functioning of both the nervous and immune systems.

By contrast, and because of the nature of neuropeptides, the analysis of *TRP* and *TRPR* gene expression was enriched by examining the relative abundance of TRP precursor in the *T. molitor* nervous system during immune system activation. Generally, neuropeptides are synthesized and stored in neuroendocrine cells, from which they are released in response to specific physiological triggers [17]. Our study revealed that the significant decrease in the expression level of *TRPR* is associated with a lower abundance of TRP expressed as an immunofluorescent signal in the nervous system, especially at the initial stage of immune system activation. This observation may suggest that TRP can be released into the haemolymph. Thus, we cannot exclude the possibility that the significant downregulation of *TRPR* at this time point is a possible protective mechanism against overstimulation [18]. Similar regulations occur in vertebrates. For example, increased levels of insulin in the bloodstream lead to a decrease in the number of associated receptors. This mechanism helps maintain homeostasis and prevents detrimental effects from excessive receptor activation, paralleling how insects may modulate *TRPR* expression to manage immune responses effectively [19]. This hypothesis is also supported by the fact that the downregulation of *TRPR* is observed not only in the nervous system but also in the fat body and haemocytes. Nevertheless, in the brain, the changes in TRP abundance are less pronounced, potentially due to the role of TRP as a neuromodulator [20].

According to our hypothesis, different immune activators seem to be a key variable for TRP pathway modulation. Here, we found differences in the changes in *TRP* and *TRPR* expression levels, as well as in the relative abundance of TRP in the nervous system after the application of different bacterial components and *Tm*Spz-like cytokine. In general, the application of *Ec* and *Tm*Spz-like elicited more significant changes in the expression levels of genes related to TRP signalling and potential TRP abundance in the nervous system than PG did. Furthermore, *Ec* significantly decreased the survival rate of the tested *T. molitor*. These differences may suggest a relationship between TRP signalling and the Imd pathway, but also Toll pathway, in which the *Tm*Spz-like cytokine is an important component [21]. *Tm*Spz-like proteins play a key role in the immune response. They are known to interact with Toll receptors, initiating immune system signalling cascades that enhance defense mechanisms. *Tm*Spz-like proteins appear later in the immune response, which may classify them as late response indicators [21]. Interestingly, our previous research also suggested that *Tm*Spz-like action can be linked to the regulation of the *T. molitor* neuroendocrine system [12]. In vertebrates, different cytokines regulate innate and adaptive immunity, activate inflammatory responses, and direct leukocyte trafficking [22]. These inflammatory factors can also affect the expression of genes related to the function of the neuroendocrine system. For example, when cytokines are present, the expression of SP is significantly increased [16]. Our results seem to confirm the existence of similar mechanisms in insects. Three hours after *Tm*Spz-like protein injection, a noticeable decrease in the abundance of the immunofluorescent signal for TRP in the VNC was observed. To our knowledge, this is the first evidence that insect cytokines may affect the abundance of neuropeptides in insect neuroendocrine cells and probably also lead to neuropeptide release to the haemocoel. Nevertheless, a significant increase in *TRP* expression in the VNC was observed at this time point. This observation can be explained by a compensatory response to the initial depletion of the TRP precursor pool in neuroendocrine cells. These findings suggest that *Tm*Spz-like is not only a regulator of immune system function but also a neuroendocrine regulator. However, to fully confirm this hypothesis, further research is needed.

The potential relationships between TRP signalling and immune system activity are also apparent in the correlation of the expression levels of *TRP, TRPR*, and immune-related genes in fat body. For a better understating role of TRP signalling and insect immune system the most important seems to be a significant positive correlation between the expression level of *TRPR* and the expression level of *Toll* and *Domeless*. Since TRPs can regulate processes related to immunomodulation and can also act as stress mediators, there may be coregulation of the expression of their receptors and those responsible for the immune response. This crosstalk between signalling pathways can help respond effectively to changing environmental conditions [15,23]. Pearson correlation coefficient analysis also revealed very interesting dependencies in the *Tenebrio* fat body because the significant correlation of the expression of the *Toll* gene in the fat body was linked to the expression of the *TRPR* gene in the brain, VNC, and fat body. The results of further experiments seem to confirm this close connection between TRP and the Toll pathway because injection of the TRPR agonist Spantide II elicits the opposite effect (upregulation) on the expression level of the *Toll* gene, as does the application of Tenmo-TRP-7 (downregulation). Importantly, simultaneous injection of Tenmo-TRP-7 and Spantide II did not affect the expression level of the *Toll* gene. On the other hand, opposite results were observed after dsRNA treatment. Simultaneous knockdown of *TRP* and *TRPR* led to significant downregulation of the *Toll* gene. The differences between these experiments may be associated with the different impacts of these methods on the TRP system. By blocking TRP receptors, Spantide II allows us to assess how the inhibition of TRP signalling affects the immune response. On the other hand, dsRNA-mediated gene knockdown targets specific genes involved in TRP pathways, providing insights into how the downregulation of these genes impacts gene expression patterns and immune signalling. However, dsRNA treatment did not completely inhibit TRP signalling because, owing to the nature of neuropeptides, some portions of TRPs may be present in the neuroendocrine system, and the knockdown of *TRPR* did not exclude the presence of active TRPR on the cell surface. For this reason, the use of Spantide II and dsRNA enables the observation of both physiological and molecular changes, offering a comprehensive approach to studying the role of TRP signalling in immune processes.

The interaction between TRP signalling and Toll pathways in the fat body cannot be considered only in terms of immune system activity but also in terms of insect metabolism [24]. Activation of the Toll pathway in the fat body, which has a metabolic-immune function similar to that of adipose tissue in mammals, can lead to a widespread metabolic switch during infection [24]. The main mechanism behind this switch is the suppression of insulin-like peptide (ILP) signalling, resulting in a reduction in triglyceride stores and a global inhibition of body growth [25,26]. Additionally, recent studies in *Drosophila* also indicate that TRPs inhibit protein-rich food intake and increase sugar appetite which is mediated by glucagon-like adipokinetic hormone (AKH) signalling [27,28]. Interestingly, all of these neuropeptides (ILPs and AKHs) possess confirmed immunomodulatory properties [29,30]. Moreover, research by Fusca et al. [31] on the cockroach *Periplaneta americana* revealed the colocalization of neurons associated with allatotropins (ATs) and TRP. Our previous research showed that AT has immunomodulatory properties in *T. molitor*. Pearson correlation analysis of available data published by Konopińska et al. [12] revealed significant links between components of the AT and TRP systems, suggesting their coordinated role in regulating immune and metabolic responses during infection (Fig. S8). For these reasons, we cannot exclude the possibility that the observed effects are the result of direct immunomodulatory action of TRP and other neuropeptides. Nevertheless, all the mentioned results point to a close connection between the TRP system and the Toll pathway in *T. molitor*.

The activation of the immune system also revealed a significant correlation between TRP system elements and the expression of *Domeless* genes in fat body, which are key components of the JAK/STAT pathway. This pathway regulates immune responses and metabolism during infection. In *Drosophila*, cytokines activate JAK/STAT via *Domeless* in tissues such as muscles, promoting immune functions and reallocating energy by suppressing ILPs, lowering glycogen, and increasing glucose [32]. The correlation between *Domeless* expression in the fat body and *TRP/TRPR* in nervous tissue suggests cooperation in immune and metabolic regulation. However, Spantide II and dsRNA treatment only partially confirmed the direct dependencies of the TRP system on JAK/STAT in the fat body. The injection of Tenmo-TRP-7 significantly downregulated the *Domeless* gene, and the simultaneous injection of Tenmo-TRP-7 with Spantide II abolished this effect. Interestingly, the application of dsControl and the administration of Tenmo-TRP-7 had effects similar to those of Tenmo-TRP-7 alone, whereas the knockdown of the expression of the *TRPR* gene and Tenmo-TRP-7 injection caused the overexpression of *Domeless*. This result may indicate that TRP-related signalling is effectively blocked. However, after double knockdown, significant changes in *Domeless* expression have not been reported.

Only weak correlations were found between TRP signalling genes and *Relish*, a key Imd pathway gene, in the fat body during immune activation in *T. molitor*. Kamareddine et al. [33] showed that the Imd pathway can be involved in the regulation of host metabolism during infection, probably *via* TRP signalling. Despite the small number of correlations between genes associated with TRR signalling and *Relish*, Spantide II treatment and dsRNA application led to interesting results. Tenmo-TRP-7, Spantide II, or their combination did not affect *Relish*, but all the treatments downregulated *Attacin 2*, an AMP mostly linked to Imd pathway [34]. Double knockdown of *TRP* and *TRPR* decreased *Relish* expression without affecting *Attacin 2*, whereas the combination of Tenmo-TRP-7 and ds*TRPR* led to *Relish* overexpression. These results can be related to the newest finding which indicate that the Attacins synthesis may be also co-regulated via Toll signalling, which is strongly affected by TRP [13]. To gain a deeper understanding of the relationship between TRP signalling and Imd pathways in the fat body, further research is needed.

Conducting analyses not only on the fat body but also on haemocytes confirms that TRP signalling may be crucial for modulating the activity of these cells. This is indicated, among other things, by a significant correlation between genes associated with TRP signalling and key immune genes. Similar to fat body, a positive correlation was observed between the expression of *TRPR* in haemocytes and the expression of *Toll* and *Domeless*, in these cells. However, the positive correlation was observed also in case of expression level *TRPR* and *Tenecin 3*. The strong link between TRP receptors and the expression of immune-related genes in haemocytes suggests that TRP signalling may be directly involved in modulating immune signalling pathways in haemocytes. These results support our previous findings because the application of Tenmo-TRP-7 significantly affects haemocyte morphology [4]. Additionally, the effects of Spantide II and dsRNA treatments further confirmed the importance of TRP signalling in haemocyte function. The application of Spantide II and blocking TRP receptor led to the upregulation of all immune-related genes in haemocytes. Moreover, a mixture of Tenmo-TRP-7 and a TRPR antagonist abolished these effects, which point to direct effect of Tenmo-TRP-7 on the expression of tested genes in haemocytes. Interestingly, only simultaneous knockdown of *TRP* and *TRPR* evoked effects such as those observed in the case of Spantide II, and significant upregulation of the genes encoding Attacin 2 and Tenecin 3 was reported. However, in the variants subjected to double injection (knockdown + physiological saline or Tenmo-TRP-7), *Attacin 2* and *Tenecin 3* were also significantly upregulated. In these cases, we cannot exclude the possibility that this can be an effect of repeated mechanical injury associated with the injection. The relationship between TRP signalling and *Tenebrio* haemocyte activity can be supported by the results of the analysis of PO activity, the enzyme produced and secreted by haemocytes in proenzyme form (prophenoloxidase, proPO). Silencing of the gene encoding the TRP precursor and simultaneous silencing of *TRP* and *TRPR* expression resulted in the inhibition of PO activity. However, in some cases, an increase in PO activity was reported, for example, after the receptor was suppressed through Spantide II and after the mixture of Spantide II and Tenmo-TRP-7 was used. These findings are consistent with transcriptomic data indicating that TRP signalling may differentially modulate the expression of PO-related genes [15]. These results suggest a complex, potentially dual role of TRP signalling in regulating PO activity, encompassing both immune-related functions and cuticular melanization processes, which may be mediated by the involvement of TRP in maintaining water balance [35–37]. Moreover, similar to the fat body, TRP signalling action on haemocytes also may be directly related to the modulation of haemocyte metabolism, which will be the subject of our future research.

Despite the undeniable influence of the TRP system on *T. molitor* immunity, from a physiological perspective, the most critical question is how TRP modulation affects the lifespan of the tested beetles. Results of the survival experiments further underscore the complexity and multidimensional nature of this issue. While Tenmo-TRP-7 and Spantide II alone had no effect, their combination reduced *T. molitor* lifespan. What is important, *TRP* knockdown decreased mortality, suggesting that partial inhibition of the TRP system might support longevity probably by modulating stress response, immunity, and metabolism. However, this effect was not detected with *TRPR* or combined *TRP/TRPR* knockdown, which requires further study. Additionally, considering that *Ec* activates the Imd and Toll pathway, particularly triggering antimicrobial peptide expression, and reduce the beetle survival it is important to assess whether the modulation of TRP signalling has any measurable effect on insect survival under such immune-challenging conditions. Gehan-Breslow-Wilcoxon analyses revealed no significant differences between groups after TRP, Spantide II, or double-knockdown application and immune system activation via *Ec*. These results may suggest that the effect of TRP is subtle or compensated for by the activation of signalling pathways related to other neuropeptides. However, strong suppression of TRP signalling by double-knockdown of *TRP* and *TRPR* reduce *T. molitor* mortality after *Ec* injection, which is a significant confirmation close relationship between TRP signalling and *T. molitor* immune system. Our supposition support also fact that, injection of Tenmo-TRP-7 reduced beetle survival ratio, especially during the first days after *Ec* administration.

Our results indicate that the TRP system in *T. molitor* plays an immunomodulatory role. We demonstrate that both *TRP* and *TRPR* expression are closely correlated with key immune-related genes, particularly those involved in the Toll and JAK/STAT pathways. Functional assays using Spantide II and RNAi-mediated gene silencing further confirm that TRP signalling can modulate immune gene expression in fat body and haemocytes, and the activity of phenoloxidase. Importantly, we show for the first time that TRP signalling is influenced by insect cytokines such as *Tm*Spz-like. Although some effects may be secondary and linked to roles in energy mobilization and stress responses through other neuropeptides which highlight the need for further studies. These investigations are important because they not only increase our understanding of insect physiology but also have practical implications for agriculture and pest management. By elucidating the roles of TRPs in immune and metabolic processes, we may identify novel targets for pest control strategies that exploit these pathways, potentially leading to more sustainable and effective approaches to managing populations of pest insects. On the other hand, some variants significantly enhance the expression of AMP genes and reduce beetle mortality, which is desirable for optimizing insect mass-rearing processes.

## 4. Materials and methods

### 4.1. Insects

The model organisms used in the experiment were 7–8-day-old adult males of *Tenebrio molitor* to avoid effects related to aging of the immune system and hormonal changes associated with oogenesis in females. Insects were reared at the Department of Animal Physiology and Developmental Biology (Adam Mickiewicz University, Poznań, Poland) in MIR 154-PE incubators (PHCbi, Singapore, Republic of Singapore) in full darkness at 28°C and 50–60% humidity. The larvae were kept in boxes with oatmeal, to which apple slices were added 3 times a week, and the pupae were removed. The adult insects used in the experiment were kept in sterile dishes with oatmeal and a piece of apple.

### 4.2. Injection and sample collection

Before the injection of the test compounds and sample collection, the insects were anaesthetized for 7 min with endogenous carbon dioxide and then washed in ethanol and distilled water. Then, with a microliter syringe (Hamilton Company, Reno, NV, USA), individuals were injected under the coxa of the third pair of legs with 2 µL of the tested solutions. Since *T. molitor* commonly experiences injuries and the injection of sterile saline has minimal impact [38], the study did not include a control group of unmanipulated males.

Selected insect tissues were collected *via* a Zeiss Stemi 504 microscope (Zeiss, Jena, Germany) under sterile conditions. To collect the haemolymph with haemocytes, the tibia of the first pair of legs was cut to obtain a sample without contamination from other cells [12].

### 4.3. Activation of *T. molitor* immune system

The immune system activation procedure was performed according to a method previously described by Konopińska et al. [12]. To activate different signalling pathways involved in immune system activation, insects were injected with 2 μL of a suspension of lyophilized *E. coli* K12 (*Ec*, 1 mg/mL, OD_600_ = 1; Sigma Aldrich, Saint Louis, MO, USA) or peptidoglycan of *S. aureus* (PG, 1 mg/mL, OD_600_ = 1; Sigma Aldrich, Saint Louis, MO, USA) in physiological saline (PS, 128 mM NaCl, 18 mM CaCl_2_, 1.3 mM KCl, 2.3 mM NaHCO_3_). Moreover, to evaluate the effects of cytokines, the *Tm*Spz-like protein was also used. The protein was synthesized by Biomatik (Kitchener, Canada) on the basis of available sequences [14] and transcriptomic data (https://www.ncbi.nlm.nih.gov). Owing to the EC_50_ value, the TmSpz-like solution in PS was injected at a concentration of 10^− 7^ M, which effectively activated the Toll pathway while avoiding toxicity and nonspecific effects [39]. As the total volume of *Tenebrio* haemolymph is approximately 18 µL, the final concentration in the insect’s body was 10^−8^ M [40]. Controls were injected with PS. After 3, 6, or 24 hours, the brain, VNC, haemolymph, and fat body were isolated. The time variants were selected on the basis of previous studies and the available literature [12,13].

#### 4.3.1. Survival experiment

The survival experiments were performed as described by Urbański et al. [15] After the injection of immune activators, 10 males were kept in a plastic box for 21 days. The beetles were kept under the same conditions as the other individuals used in this study. The number of dead and living individuals was checked every day during the experiment. Dead individuals were removed from the boxes. Ten individuals kept in plastic boxes were considered one biological replicate. Each research variant was repeated at least five times (5 × 10 individuals = 50 beetles per treatment).

#### 4.3.2. Expression levels of tested genes after activation of the *T. molitor* immune system

At selected time points after activation of the *Tenebrio* immune system, the brain, VNC, haemolymph, and fat body were isolated from each individual and transferred to separate Eppendorf tubes containing 200 µL of lysis buffer (Zymo Research, Irvine, CA, USA). The collected samples were frozen in liquid nitrogen and stored at -80°C. Before RNA isolation, the samples were homogenized for 1 min *via* a pellet homogenizer (Kimble Chase, Vineland, NJ, USA). RNA isolation was performed with a Quick-RNA Mini-prep kit (Zymo Research, Irvine, CA, USA) according to the manufacturer’s protocol. To eliminate DNA residues, the samples were subsequently incubated with a Turbo DNA-free kit (Thermo Fisher Scientific, Waltham, MA, USA). The quality and quantity of the obtained RNA were analysed with a DS-11 spectrophotometer (DeNovix, Inc., Wilmington, DE, USA).

cDNA was synthesized from equal concentrations of RNA templates (100 ng for brain, VNC, fat body, and 50 ng for haemocytes) *via* a LunaScript® RT SuperMix Kit (New England Biolabs, Ipswich, MA, USA). Additional quality control (no RT control) was also performed. For RT‒qPCR analysis, the expression levels of genes encoding TRP precursor and TRPR were analysed. Moreover, in both experiments, the expression levels of selected immune-related genes were estimated. The primers used in the experiment were synthesized by the Institute of Biochemistry and Biophysics of the Polish Academy of Science (Warsaw, Poland). To confirm our results, the amplicons were sequenced by the Molecular Biology Techniques Laboratory (Faculty of Biology, Adam Mickiewicz University, Poznań, Poland) and compared with data available in a public database (https://www.ncbi.nlm.nih.gov). The expression of selected genes was normalized on the basis of the expression level of the gene encoding *T. molitor* ribosomal protein L13a (*TmRpL13a*) [41]. The sequences of the primers used are available in Table S1. The efficiency of the primers used ranged from 92–108%. RT‒qPCR analysis was performed on a QuantStudio™ 3 Real-Time PCR System, 96-well, 0.2 mL (Applied Biosystems™, Waltham, MA, USA) and a Corbett Research RG-6000 Real-Time PCR Thermocycler (Qiagen, Hilden, Germany). Three biological replicates and at least two technical repeats were used for each repeat. In one biological replication, tissues from at least 20 individuals (for the brain, VNC, and haemolymph) or 10 individuals (for the fat body, owing to its specific structure) were collected for each sample. The relative expression was calculated *via* the method described by Plaffl [42].

#### 4.3.3. Immunolocalization of TRP precursor in the nervous system after activation of the immune system

Analysis of changes in the distribution and abundance of the TRP precursor after immune system activation was performed according to the method described by Urbański et al. [43]. The immune system of the insects was activated as described in section 2.4. The control was PS-injected individuals. After 3, 6, and 24 h, the brain and VNC were isolated. The isolated structures were left in 2% paraformaldehyde for 24 h. Then, the samples were washed (4 × 1 h) in phosphate-buffered saline (PBS) and incubated overnight at 4°C in a solution containing 4% Triton-X 100 (PBSTX) (Sigma‒Aldrich, St. Louis, MO, USA), 2% normal goat serum (NGS, Jackson ImmunoResearch Lab., West Grove, PA, USA) and 2% bovine serum albumin (BSA, Sigma‒Aldrich, St. Louis, MO, USA). Next, they were incubated with primary antibodies (1:500, anti-Aedae-TRP-2, gifted from Prof. J. Veenstra) [44] with 2% NGS, 2% BSA, and 0.4% Triton-X at 4°C for 72 hours. The samples were washed again in PBSTX (4 × 1 h), and then secondary antibodies in PBSTX (1:200) were added and incubated for 24 h at 4°C in the dark. The samples were washed with PBS (4 × 15 min) and sealed on a microscope slide in 90% glycerol with DABCO. In addition, two negative controls were made, one without the primary antibodies and one without secondary antibodies. At least two biological replicates were performed for each variant. The samples were analysed with a confocal microscope (LSM 510, Axiovert 200 M, Carl Zeiss, Oberkochen, Germany). The samples related to the selected time points were collected at the same time, including the control individuals. Additionally, sample examination was performed on the same day under the same conditions and microscope setup, which allowed comparison of the obtained micrographs. For a detailed analysis of the tested parts of the *Tenebrio* nervous system, each of the structures was examined *via* a Z-stack module. Z-stack files based on the fluorescence intensity (threshold 75–255) were used for 3D visualization of the distribution and abundance of fluorescence-positive signals related to the presence of the TRP precursor. 3D visualization was performed with AMIRA 3D software (Thermo Fisher Scientific, Waltham, MA, USA).

### 4.4. Effects of Spantide II and dsRNA on selected immune mechanisms of *T. molitor*

#### 4.4.1. Antagonist of TRPR – Spantide II

Spantide II (Sigma‒Aldrich, St. Louis, MO, USA) was chosen for its higher affinity for NK1 and insect TRPR than Spantide I and Spantide III [45]. On the basis of the literature data, in the experiment, insects were injected with a solution of Spantide II in PS at a concentration of 10^-3^ M. On the basis of the literature concerning the total haemolymph volume of adult *T. molitor* (approx. 18 μL), the final concentration of Spantide II was 10^-4^ M [40,46]. In the experiment with Spantide II, the effect of Tenmo-TRP-7 (amino acid sequence: MPRQSGFFGMRa) on selected immune bioassays was also tested as an internal control of the effect of TRP signalling activation. Tenmo-TRP-7 was synthesized by Creative Peptides (Shirley, NY, USA; purity >95% HPLC) and was used at concentrations of 10^-5^ M in PS. The concentrations were chosen on the basis of literature data and our previous studies [4,10]. Moreover, simultaneous injection of Tenmo-TRP-7 and Spantide II was also evaluated. Previously, the strongest immunomodulatory effect of Tenmo-TRP was observed 24 hours after its injection. For this reason, the effects of Spantide II and Tenmo-TRP-7 were evaluated at this time point [4,10]. The negative control was injected with PS.

#### 4.4.2. Synthesis of specific dsRNA-targeted genes encoding TRP precursor and receptor

dsRNAs directed against the precursor and TRP receptor were synthesized via a modified method described by Zanchi et al. and Keshavarz et al. [47,48]. The first step of specific dsRNA synthesis was the extraction of total RNA from adult *T. molitor* and larvae of *Galleria mellonella* (control) with the Quick-RNA Mini-prep Kit (Zymo Research, Irvine, CA, USA). As an internal control, dsRNA based on *G. mellonella* lysozyme cDNA, which has no sequence homology with any known gene of *T. molitor*, was used. The prepared samples were subsequently incubated with a TurboDNA-free kit (Thermo Fisher Scientific, Waltham, MA, USA). The RNA concentrations and quality were checked with a DS-11 spectrophotometer (DeNovix, Inc., Wilmington, DE, USA). The cDNA was subsequently synthesized on an RNA template via the LunaScript® RT SuperMix Kit (New England Biolabs, Ipswich, MA, USA), and the quality of the RNA was checked *via* a no-RT control. The fragments were amplified via PCR with the KAPA2G Fast ReadyMix PCR Kit (KAPA Biosystems, Sigma‒Aldrich, St. Louis, MO, USA) via a template derived from synthesized cDNA. The primer sequences are available in Table 1. To check the product quality, the obtained amplicons were analysed via electrophoresis via a 2% TAE agarose gel with ethidium bromide. Next, the samples were cleaned with the PCR/DNA Clean-Up DNA Kit (EURx, Gdańsk, Poland), and dsRNA synthesis was initiated via the HighYield T7 RNA Synthesis Kit (Jena Bioscience GmbH, Jena, Germany) according to the manufacturer’s protocol. The incubation time at 37°C was extended to 4 hours. To remove DNA residues, the samples were additionally incubated with a TurboDNA-free kit. For RNA hybridization, the samples were heated to 95°C and left in a thermocycler to cool overnight. The synthesized dsRNA was then washed with a 5 M NH_4_OAc (Thermo Fisher Scientific, Waltham, MA, USA) and ethanol gradient (Bioultra, molecular grade) (70%-99%). The dsRNA pellet was then resuspended in nuclease-free water and kept at -20°C until further use. For the suppression of the targeted genes, beetles were injected with 2 μg of prepared dsRNA (1 μg/μL in 2 μL of water). For double-knockdown, beetles were injected with 4 μg of prepared dsRNA (2 μg/μL of dsRNA *GmLys* in 2 μL of water) as a control or with 4 μg of a mixture of dsRNA-targeted genes encoding TRP precursor (2 μg) and receptor (2 μg).

Before the experiments, the knockdown efficiency was estimated by extraction of total RNA (n=3, pools of three adults per day) at different time points after exposure (Fig. S9). The samples were then processed as described above. The relative expression was evaluated via RT‒qPCR as described in section 2.2.1. On the basis of the obtained results, the day on which the studied genes were silenced was selected.

For the double-injection variant, individuals were injected with dsRNA directed against the gene encoding *Galleria* lysozyme or TRPR. After 8 days, the insects were injected with physiological saline or Tenmo-TRP-7 at a concentration of 10^-5^ M. The control group was injected with *dsGmLys* and physiological saline.

#### 4.4.3. Survival experiment after injection of Tenmo-TRP-7, Spantide II or dsRNA

A survival experiment involving the different variants with Tenmo-TRP-7, Spantide II and dsRNA treatment was performed according to the methods described in section 4.3.1.

#### 4.4.4. The expression levels of immune-related genes after the modulation of TRP signalling

The determination of changes in the expression levels of immune-related genes after the modulation of TRP signalling via Spantide II and dsRNA in the fat body and haemocytes was performed according to the methods described in section 4.3.2.

#### 4.4.5. The activity of phenoloxidase in *T. molitor* haemolymph after the modulation of TRP signalling

Phenoloxidase activity was measured *via* a modified method published by Sorrentino et al and Urbański et al. [49,50]. A haemolymph sample (1 µL) was transferred to a paper filter (Whatman No. 52, Sigma‒Aldrich, St. Louis, MO, USA) soaked with DL-DOPA (Sigma‒Aldrich, St. Louis, MO, USA) solution (2 mg/1 mL) in 10 mM phosphate buffer. Two technical repeats were performed for each individual. At least 10 biological replicates were performed in each group. The samples were then incubated for 30 min in the dark. After drying, they were scanned with a SHARP AR 153 EN (600 dpi, 8 bits, grayscale) and analysed via ImageJ software (version 2). For each sample, a ‘mean pixel value’ was measured in its central part (40 pixels). To better visualize the changes occurring after the application of the neuropeptide under study, the results are presented as a percentage of change relative to control individuals.

### 4.5. Survival assessment following TRP system suppression and immune activation with *E. coli*

Insects were injected with physiological saline, Tenmo-TRP-7 at a concentration of 10^-5^ M, or Spantide II at a concentration of 10^-3^ M. After 2 hours, the immune system was activated. *E. coli* was chosen to trigger the immune response because of its strong impact on *T. molitor* survival. Additionally, to examine how the simultaneous silencing of *TRP* and *TRPR* affects the ability of the beetle to survive immune system activation, 8 days after dsRNA injection, the insects were injected with *E. coli*. The rest of the experiment followed the procedure described in section 4.3.1.

### 4.6. Statistical analyses

Statistical analysis was performed via GraphPad Prism 9 software (Adam Mickiewicz University licence). The outliers were identified *via* the ROUT method. The normality of the distribution was tested via the Shapiro‒Wilk test. Depending on the number of tested research variables, results consistent with the normality of the distribution were analysed by one-way ANOVA with Dunnett’s *post hoc* test or Student’s *t* test (depends on the number of analysed groups). The results with a nonnormal distribution were analysed with the Kruskal‒Wallis test with Dunn’s *post hoc* test and the Mann‒ Whitney U test (depends on the number of analysed groups). The correlation of the data was analysed via the Pearson correlation coefficient method in SRplot software (https://www.bioinformatics.com.cn/srplot). In the manuscript, only statistically significant correlations were described. Survival curves were estimated on the basis of the Kaplan‒Meier estimator. The differences between survival curves were calculated via the Gehan-Breslow-Wilcoxon test.

## Acknowledgement

This research was supported by Grant No. 2021/41/B/NZ9/01054 from the National Science Centre (Poland). Special thanks go to Prof. J. A. Veenstra for sharing the anti-TRP antibodies. We would also like to thank Prof. Guy Smagghe for his valuable comments during data interpretation, and Natalia Bylewska, Radosław Gmyrek, Karolina Kozłowska, and Sara Tchórzewska for their help in rearing *T. molitor*.

During the preparation of this work, the authors used Rubriq software (American Journal Experts (AJE)) for language correction. After using this tool, the authors reviewed and edited the content as needed and take full responsibility for the content of the publication.

## Author contributions

Conceptualization - NK, AU and JR; Sample collection – NK, AU, KWN and SC, Methodology – NK, AU, MK and GN, Performing analyses – NK, AU, KWN, SC; Visualization – NK and AU; Writing – Original Draft Preparation – NK, Writing – Review & Editing – NK, KWN, SC, GN, MK, JR and AU; Supervision – AU.

## Competing interests

The authors declare no competing interests. GN is employed by the company genXone S.A. The remaining authors declare that the research was conducted in the absence of any commercial or financial relationships that could be construed as potential conflicts of interest.

## Supporting information

**Fig. S1**. Changes in the expression levels of genes encoding TRP precursor in the brain (A-C), ventral nerve cord (D-F) and fat body (G-I) of *T. molitor* after activation of the immune system. The immune response was elicited by injection of *Escherichia coli* K16 (Ec), peptidoglycan from *Staphylococcus aureus* (PG), or Spätzle-like protein. Control – individuals injected with physiological saline. Due to the dynamic nature of the immune response, samples were collected 3, 6 and 24 hours after immune system activation.

**Fig. S2**. Changes in the expression levels of genes encoding TRP receptor (TRPR) in the brain (A-C), ventral nerve cord (D-F), fat body (G-I) and haemocytes (J-L) of *T. molitor* after activation of the immune system. The immune response was elicited by injection of *Escherichia coli* K16 (Ec), peptidoglycan from *Staphylococcus aureus* (PG), or Spätzle-like protein. Control – individuals injected with physiological saline. Due to the dynamic nature of the immune response, samples were collected 3, 6 and 24 hours after immune system activation.

**Fig. S3**. Changes in the expression levels of immune-related genes (Toll (A-C), Relish (D-F), Domeless (G-I), Attacin 2 (J-L), Tenecin 3 (M-O)) in the fat body of *T. molitor* after activation of the immune system. The immune response was elicited by injection of *Escherichia coli* K16 (Ec), peptidoglycan from *Staphylococcus aureus* (PG), or Spätzle-like protein. Control – individuals injected with physiological saline. Due to the dynamic nature of the immune response, samples were collected 3, 6 and 24 hours after immune system activation.

**Fig. S4**. Changes in the expression levels of immune-related genes (Toll (A-C), Relish (D-F), Domeless (G-I), Attacin 2 (J-L), Tenecin 3 (M-O)) in the haemocytes of *T. molitor* after activation of the immune system. The immune response was elicited by injection of *Escherichia coli* K16 (Ec), peptidoglycan from *Staphylococcus aureus* (PG), or Spätzle-like protein. Control – individuals injected with physiological saline. Due to the dynamic nature of the immune response, samples were collected 3, 6 and 24 hours after immune system activation.

**Fig. S5**. Changes in the expression levels of immune-related genes (Toll (A-C), Relish (D-F), Domeless (G-I), Attacin 2 (J-L), Tenecin 3 (M-O)) in the fat body of *T. molitor* after application of physiological saline (control), Tenmo-TRP-7 (TRP) at a concentration of 10^-5^ M, Spantide II at a concentration of 10^-3^ M, and a mixture of Tenmo-TRP-7 and Spantide II. Also after injection of dsRNA targeted genes encoding lysozyme in *Galleria mellonella* (*LysGm*, Control) or genes related to TRP signalling (genes for TRP precursor (dsRNA TRP) or receptor (dsRNA TRPR). In addition, the double-knockdown of TRP and TRPR was analyzed. Control for the double-knockdown experiment (Control 2KD) was the injection of the double dose of dsRNA-targeted *LysGm*.

**Fig. S6**. Changes in the expression levels of immune-related genes (Toll (A-C), Relish (D-F), Domeless (G-I), Attacin 2 (J-L), Tenecin 3 (M-O)) in the haemocytes of *T. molitor* after application of physiological saline (control), Tenmo-TRP-7 (TRP) at a concentration of 10^-5^ M, Spantide II at a concentration of 10^-3^ M, and a mixture of Tenmo-TRP-7 and Spantide II. Also after injection of dsRNA targeted genes encoding lysozyme in *Galleria mellonella* (*LysGm*, Control) or genes related to TRP signalling (genes for TRP precursor (dsRNA TRP) or receptor (dsRNA TRPR). In addition, the double-knockdown of TRP and TRPR was analyzed. Control for the double-knockdown experiment (Control 2KD) was the injection of the double dose of dsRNA-targeted *LysGm*.

**Fig. S7**. Correlation of expression level of *TRP* and *TRPR* in different tissues/cells during activation of *Tenebrio* immune system with the expression level of genes encoding allatotropin precursor (AT) and receptor (ATR). Red squares – significant correlations associated with the TRP system presented in this article. Blue squares – significant correlations related to AT system (data previously presented by Konopińska et al. (2024)). Green squares – significant correlations between TRP and AT systems. VNC – ventral nerve cord; FB – fat body; Hem – haemocytes. To estimate the correlation of the data, the Pearson correlation coefficient method was used. The matrix was generated using SRplot software (https://www.bioinformatics.com.cn/srplot).

## Notes

### Competing Interest Statement

The authors have declared no competing interest.

## References

1. Adamski Z, Bufo SA, Chowański S, Falabella P, Lubawy J, Marciniak P, et al. Beetles as model organisms in physiological, biomedical and environmental studies–a review. Front Physiol. 2019;10: 319.

2. Ribeiro N, Abelho M, Costa R. A review of the scientific literature for optimal conditions for mass rearing Tenebrio molitor (Coleoptera: Tenebrionidae). J Entomol Sci. 2018;53: 434–454.

3. Urbanski A, Rosinski G. Role of neuropeptides in the regulation of the insect immune system– current knowledge and perspectives. Curr Protein Pept Sci. 2018;19: 1201–1213.

4. Urbański A, Konopińska N, Lubawy J, Walkowiak-Nowicka K, Marciniak P, Rolff J. A possible role of tachykinin-related peptide on an immune system activity of mealworm beetle, Tenebrio molitor L. Dev Comp Immunol. 2021;120: 104065.

5. Ordovas-Montanes J, Rakoff-Nahoum S, Huang S, Riol-Blanco L, Barreiro O, von Andrian UH. The regulation of immunological processes by peripheral neurons in homeostasis and disease. Trends Immunol. 2015;36: 578–604.

6. Hartenstein V. The neuroendocrine system of invertebrates: a developmental and evolutionary perspective. Journal of endocrinology. 2006;190: 555–570.

7. Nässel DR, Zandawala M, Kawada T, Satake H. Tachykinins: neuropeptides that are ancient, diverse, widespread and functionally pleiotropic. Front Neurosci. 2019;13: 1262.

8. Gui S-H, Jiang H-B, Xu L, Pei Y-X, Liu X-Q, Smagghe G, et al. Role of a tachykinin-related peptide and its receptor in modulating the olfactory sensitivity in the oriental fruit fly, Bactrocera dorsalis (Hendel). Insect Biochem Mol Biol. 2017;80: 71–78.

9. Vanden Broeck J, Torfs H, Poels J, Van Poyer W, Swinnen E, Ferket K, et al. Tachykinin‐like Peptides and Their Receptors: A Review. Ann N Y Acad Sci. 1999;897: 374–387.

10. Urbański A, Konopińska N, Walkowiak-Nowicka K, Roizman D, Lubawy J, Radziej M, et al. Functional homology of tachykinin signalling: The influence of human substance P on the immune system of the mealworm beetle, Tenebrio molitor L. Dev Comp Immunol. 2023;142: 104669.

11. Mashaghi A, Marmalidou A, Tehrani M, Grace PM, Pothoulakis C, Dana R. Neuropeptide substance P and the immune response. Cellular and molecular life sciences. 2016;73: 4249–4264.

12. Konopińska N, Gmyrek R, Bylewska N, Tchórzewska S, Nowicki G, Lubawy J, et al. The allatotropin/orexin system as an example of immunomodulatory properties of neuropeptides. Insect Biochem Mol Biol. 2024;171: 104149.

13. Park S, Jo YH, Park KB, Ko HJ, Kim CE, Bae YM, et al. TmToll-7 plays a crucial role in innate immune responses against Gram-negative bacteria by regulating 5 AMP genes in Tenebrio molitor. Front Immunol. 2019;10: 310.

14. Jang HA, Patnaik BB, Ali Mohammadie Kojour M, Kim BB, Bae YM, Park KB, et al. Tm Spz-like plays a fundamental role in response to E. coli but not S. aureus or C. albican infection in Tenebrio molitor via regulation of antimicrobial peptide production. Int J Mol Sci. 2021;22: 10888.

15. Urbański A, Johnston P, Bittermann E, Keshavarz M, Paris V, Walkowiak-Nowicka K, et al. Tachykinin-related peptides modulate immune-gene expression in the mealworm beetle Tenebrio molitor L. Sci Rep. 2022;12: 17277.

16. Suvas S. Role of substance P neuropeptide in inflammation, wound healing, and tissue homeostasis. The Journal of Immunology. 2017;199: 1543–1552.

17. Bendena WG. Neuropeptide physiology in insects. Neuropeptide systems as targets for parasite and pest control. 2010; 166–191.

18. Shankaran H, Wiley HS, Resat H. Receptor downregulation and desensitization enhance the information processing ability of signalling receptors. BMC Syst Biol. 2007;1: 1–15.

19. Lewis ST, Greenway F, Tucker TR, Alexander M, Jackson LK, Hepford SA, et al. A receptor story: insulin resistance pathophysiology and physiologic insulin resensitization’s role as a treatment modality. Int J Mol Sci. 2023;24: 10927.

20. Van Loy T, Vandersmissen HP, Poels J, Van Hiel MB, Verlinden H, Broeck J Vanden. Tachykinin-related peptides and their receptors in invertebrates: a current view. Peptides (NY). 2010;31: 520–524.

21. Yu B, Sang Q, Pan G, Li C, Zhou Z. A Toll-Spätzle pathway in the immune response of Bombyx mori. Insects. 2020;11: 586.

22. Zimmerman LM, Bowden RM, Vogel LA. A vertebrate cytokine primer for eco‐immunologists. Funct Ecol. 2014;28: 1061–1073.

23. Gonzalez-Rey E, Ganea D, Delgado M. Neuropeptides: keeping the balance between pathogen immunity and immune tolerance. Curr Opin Pharmacol. 2010;10: 473–481.

24. Dolezal T, Krejcova G, Bajgar A, Nedbalova P, Strasser P. Molecular regulations of metabolism during immune response in insects. Insect Biochem Mol Biol. 2019;109: 31–42.

25. Roth SW, Bitterman MD, Birnbaum MJ, Bland ML. Innate immune signaling in Drosophila blocks insulin signaling by uncoupling PI (3, 4, 5) P3 production and Akt activation. Cell Rep. 2018;22: 2550–2556.

26. Toprak U. The role of peptide hormones in insect lipid metabolism. Front Physiol. 2020;11: 434.

27. Nässel DR. Tachykinin-related peptides in invertebrates: a review. Peptides (NY). 1999;20: 141–158.

28. Ahrentløv N, Kubrak O, Lassen M, Malita A, Koyama T, Frederiksen AS, et al. Protein-responsive gut hormone tachykinin directs food choice and impacts lifespan. Nat Metab. 2025; 1–23.

29. Černý J, Krishnan N, Hejníková M, Štěrbová H, Kodrík D. Modulation of response to braconid wasp venom by adipokinetic hormone in Drosophila melanogaster. Comparative Biochemistry and Physiology Part C: Toxicology & Pharmacology. 2024;285: 110005.

30. Chowański S, Walkowiak-Nowicka K, Winkiel M, Marciniak P, Urbański A, Pacholska-Bogalska J. Insulin-like peptides and cross-talk with other factors in the regulation of insect metabolism. Front Physiol. 2021;12: 701203.

31. Fusca D, Schachtner J, Kloppenburg P. Colocalization of allatotropin and tachykinin‐related peptides with classical transmitters in physiologically distinct subtypes of olfactory local interneurons in the cockroach (Periplaneta americana). Journal of Comparative Neurology. 2015;523: 1569–1586.

32. McMullen E, Strych L, Chodakova L, Krebs A, Dolezal T. JAK/STAT mediated insulin resistance in muscles is essential for effective immune response. Cell Communication and Signaling. 2024;22: 203.

33. Kamareddine L, Robins WP, Berkey CD, Mekalanos JJ, Watnick PI. The Drosophila immune deficiency pathway modulates enteroendocrine function and host metabolism. Cell Metab. 2018;28: 449–462.

34. Jo YH, Patnaik BB, Hwang J, Park KB, Ko HJ, Kim CE, et al. Regulation of the expression of nine antimicrobial peptide genes by Tm IMD confers resistance against Gram-negative bacteria. Sci Rep. 2019;9: 10138.

35. Marieshwari BN, Bhuvaragavan S, Sruthi K, Mullainadhan P, Janarthanan S. Insect phenoloxidase and its diverse roles: melanogenesis and beyond. Journal of Comparative Physiology B. 2023;193: 1–23.

36. Agard MA, Zandawala M, Paluzzi J-P V. Another fly diuretic hormone: tachykinins increase fluid and ion transport by adult Drosophila melanogaster Malpighian’renal’tubules. bioRxiv. 2024; 2022–2024.

37. Johard HAD, Coast GM, Mordue W, Nässel DR. Diuretic action of the peptide locustatachykinin I: cellular localisation and effects on fluid secretion in Malpighian tubules of locusts. Peptides (NY). 2003;24: 1571–1579.

38. Goerlinger A, Develay C, Balourdet A, Rigaud T, Moret Y. Infection risk by oral contamination does not induce immune priming in the mealworm beetle (Tenebrio molitor) but triggers behavioral and physiological responses. Front Immunol. 2024;15: 1354046.

39. Weber ANR, Tauszig-Delamasure S, Hoffmann JA, Lelièvre E, Gascan H, Ray KP, et al. Binding of the Drosophila cytokine Spätzle to Toll is direct and establishes signaling. Nat Immunol. 2003;4: 794–800.

40. Walkowiak-Nowicka K, Chowański S, Pacholska-Bogalska J, Adamski Z, Kuczer M, Rosiński G. Antheraea peptide and its analog: Their influence on the maturation of the reproductive system, embryogenesis, and early larval development in Tenebrio molitor L. beetle. PLoS One. 2022;17: e0278473.

41. Jacobs CGC, Gallagher JD, Evison SEF, Heckel DG, Vilcinskas A, Vogel H. Endogenous egg immune defenses in the yellow mealworm beetle (Tenebrio molitor). Dev Comp Immunol. 2017;70: 1– 8.

42. Pfaffl MW. A new mathematical model for relative quantification in real-time RT–PCR. Nucleic Acids Res. 2001;29: e45–e45.

43. Urbański A, Lubawy J, Marciniak P, Rosiński G. Myotropic activity and immunolocalization of selected neuropeptides of the burying beetle Nicrophorus vespilloides (Coleoptera: Silphidae). Insect Sci. 2019;26: 656–670.

44. Veenstra JA, Lau GW, Agricola H-J, Petzel DH. Immunohistological localization of regulatory peptides in the midgut of the female mosquito Aedes aegypti. Histochem Cell Biol. 1995;104: 337–347.

45. Janecka A, Poels J, Fichna J, Studzian K, Broeck J Vanden. Comparison of antagonist activity of spantide family at human neurokinin receptors measured by aequorin luminescence-based functional calcium assay. Regul Pept. 2005;131: 23–28.

46. Poels J, Birse RT, Nachman RJ, Fichna J, Janecka A, Broeck J Vanden, et al. Characterization and distribution of NKD, a receptor for Drosophila tachykinin-related peptide 6. Peptides (NY). 2009;30: 545–556.

47. Zanchi C, Johnston PR, Rolff J. Evolution of defence cocktails: Antimicrobial peptide combinations reduce mortality and persistent infection. Mol Ecol. 2017;26: 5334–5343.

48. Keshavarz M, Zanchi C, Rolff J. The effect of combined knockdowns of Attacins on survival and bacterial load in Tenebrio molitor. Front Immunol. 2023;14: 1140627.

49. Sorrentino RP, Small CN, Govind S. Quantitative analysis of phenol oxidase activity in insect hemolymph. Biotechniques. 2002;32: 815–823.

50. Urbański A, Czarniewska E, Baraniak E, Rosiński G. Developmental changes in cellular and humoral responses of the burying beetle Nicrophorus vespilloides (Coleoptera, Silphidae). J Insect Physiol. 2014;60: 98–103.

