## Supplementary materials for "Tachykinin-related peptide signalling is important for the immune response of the mealworm beetle *Tenebrio molitor* L"

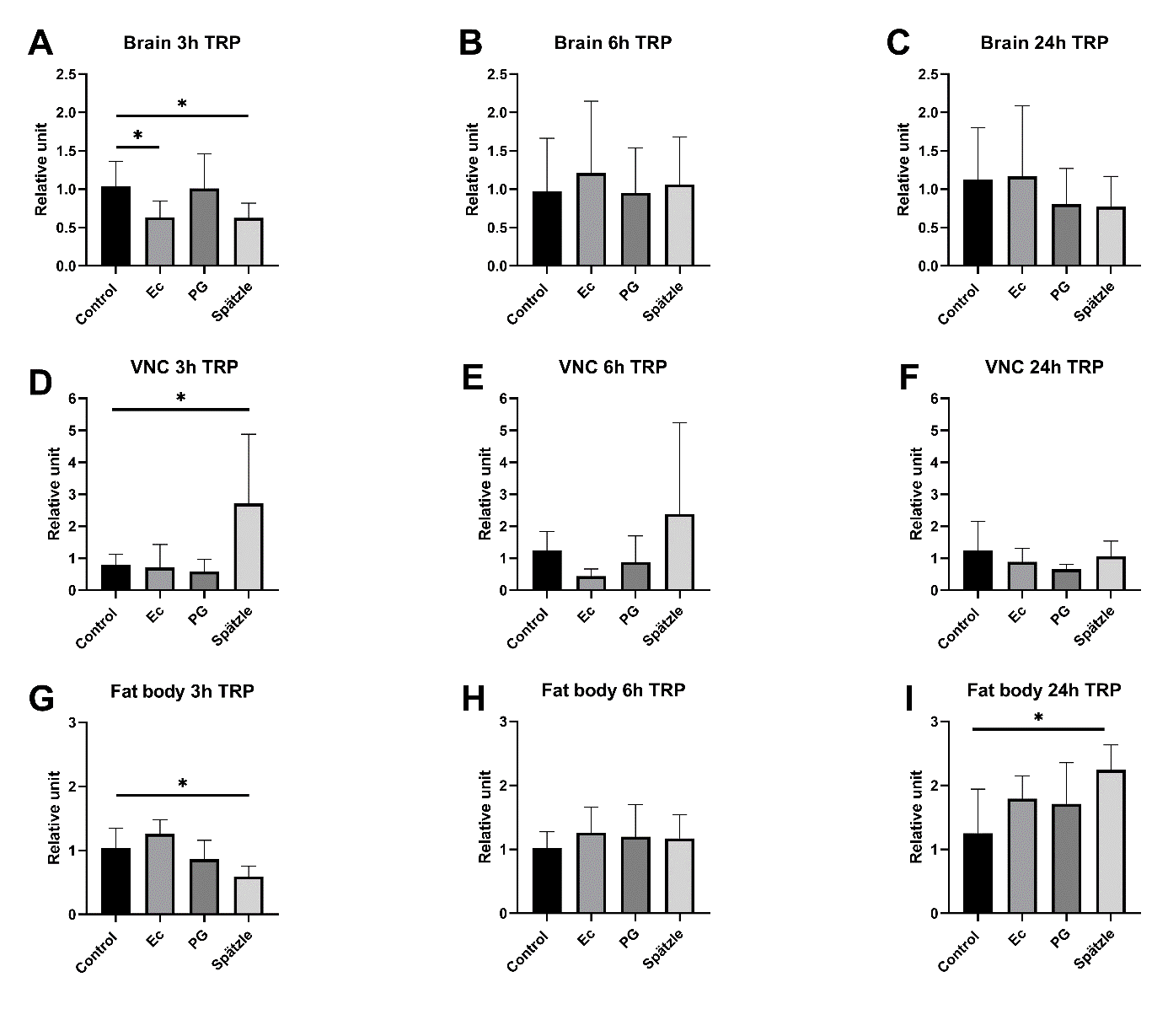

**Fig. S1.** Changes in the expression levels of genes encoding TRP precursor in the brain (A-C), ventral nerve cord (D-F) and fat body (G-I) of *T. molitor* after activation of the immune system. The immune response was elicited by injection of *Escherichia coli* K16 (Ec), peptidoglycan from *Staphylococcus aureus* (PG), or Spätzle-like protein. Control – individuals injected with physiological saline. Due to the dynamic nature of the immune response, samples were collected 3, 6 and 24 hours after immune system activation. The values are the means ± SDs. Asterisks indicate statistically significant changes compared to those in control individuals; **p*≤0.05.

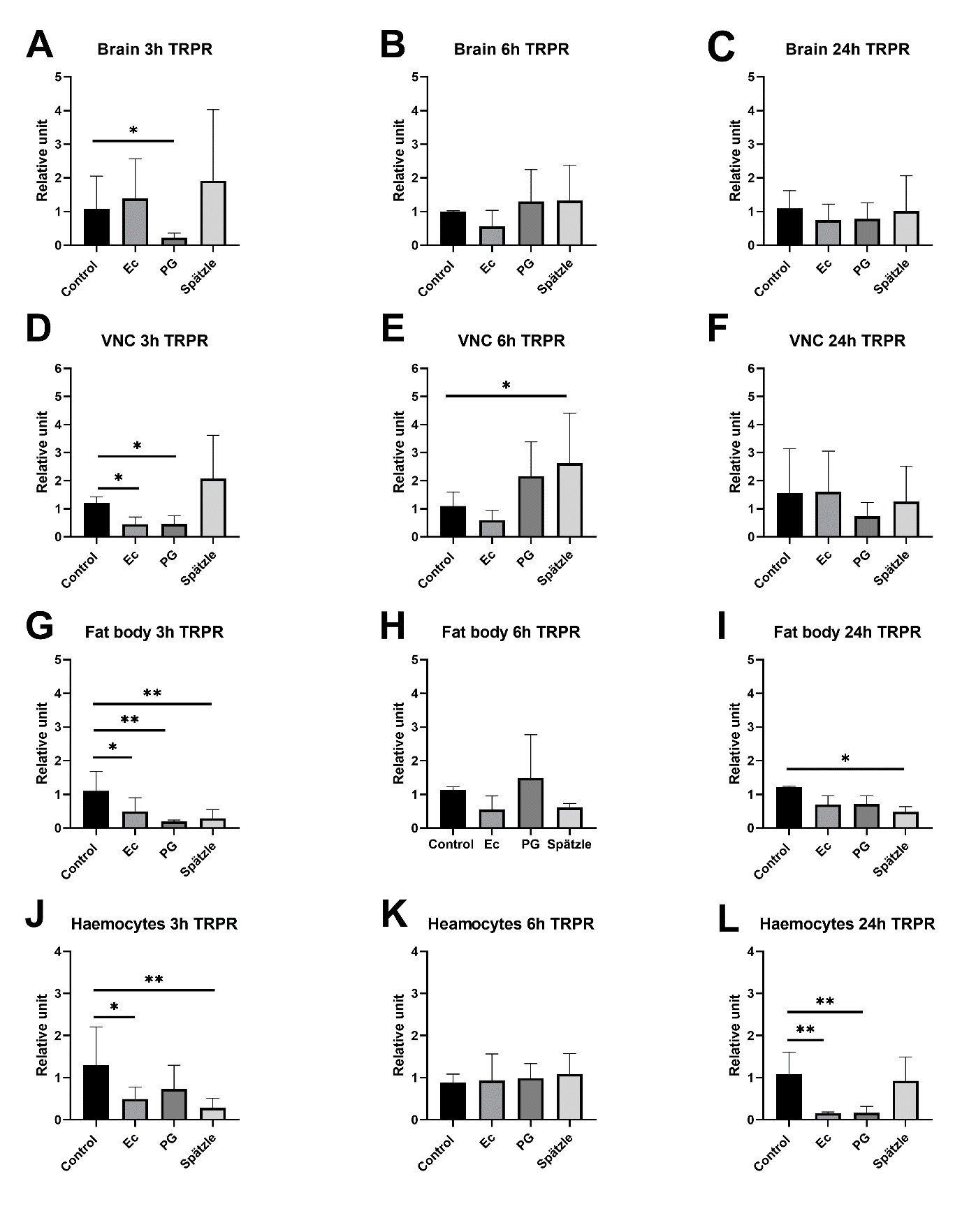

**Fig. S2.** Changes in the expression levels of genes encoding TRP receptor (TRPR) in the brain (A-C), ventral nerve cord (D-F), fat body (G-I) and haemocytes (J-L) of *T. molitor* after activation of the immune system. The immune response was elicited by injection of *Escherichia coli* K16 (Ec), peptidoglycan from *Staphylococcus aureus* (PG), or Spätzle-like protein. Control – individuals injected with physiological saline. Due to the dynamic nature of the immune response, samples were collected 3, 6 and 24 hours after immune system activation. The values are the means ± SDs. Asterisks indicate statistically significant changes compared to those in control individuals; **p*≤0.05, ***p*≤0.01.

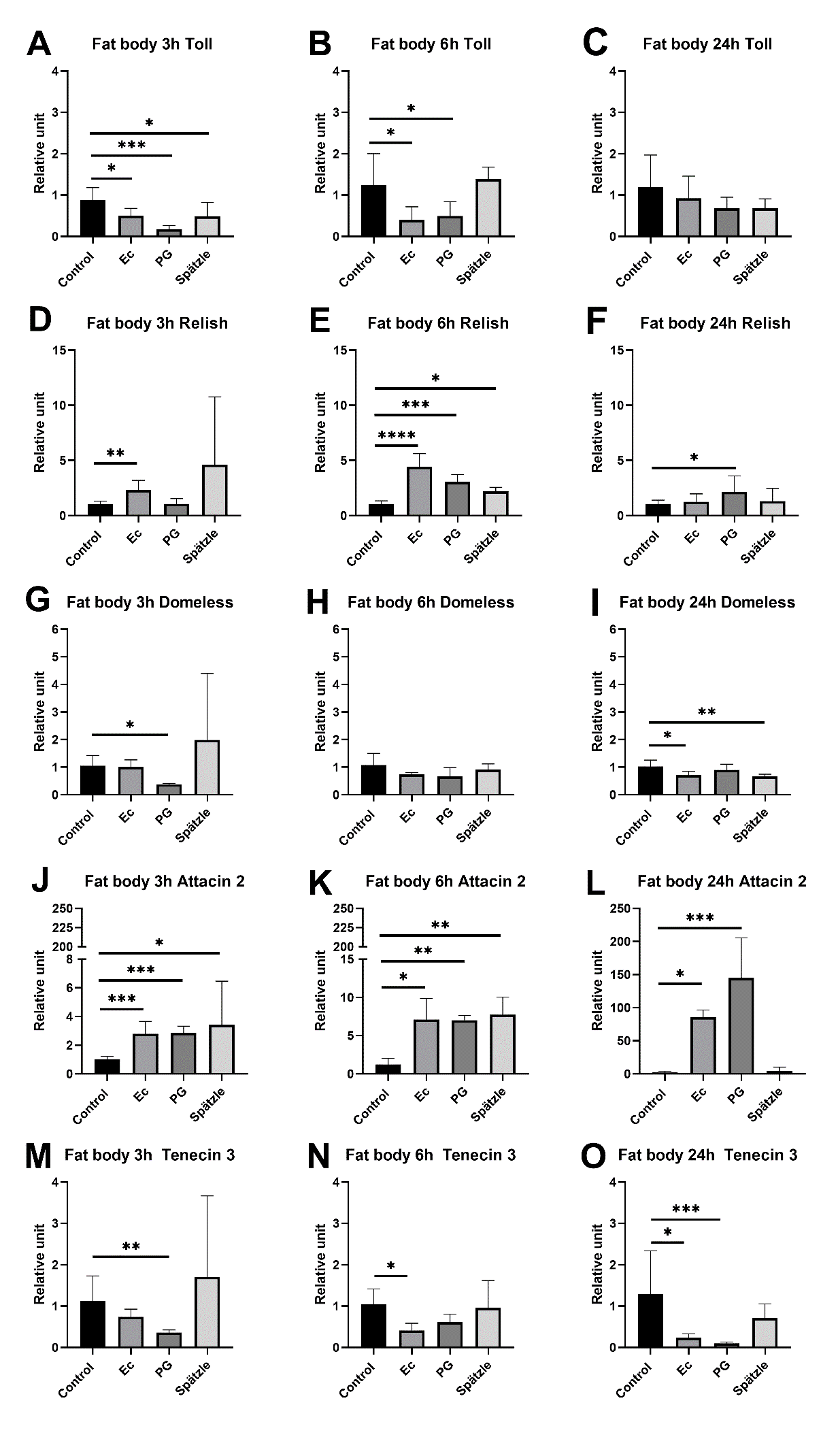

**Fig. S3.** Changes in the expression levels of immune-related genes (Toll (A-C), Relish (D-F), Domeless (G-I), Attacin 2 (J-L), Tenecin 3 (M-O)) in the fat body of *T. molitor* after activation of the immune system. The immune response was elicited by injection of *Escherichia coli* K16 (Ec), peptidoglycan from *Staphylococcus aureus* (PG), or Spätzle-like protein. Control – individuals injected with physiological saline. Due to the dynamic nature of the immune response, samples were collected 3, 6 and 24 hours after immune system activation. The values are the means ± SDs. Asterisks indicate statistically significant changes compared to those in control individuals; **p*≤0.05, ***p*≤0.01, ****p*≤0.001, **** *p*≤0.0001.

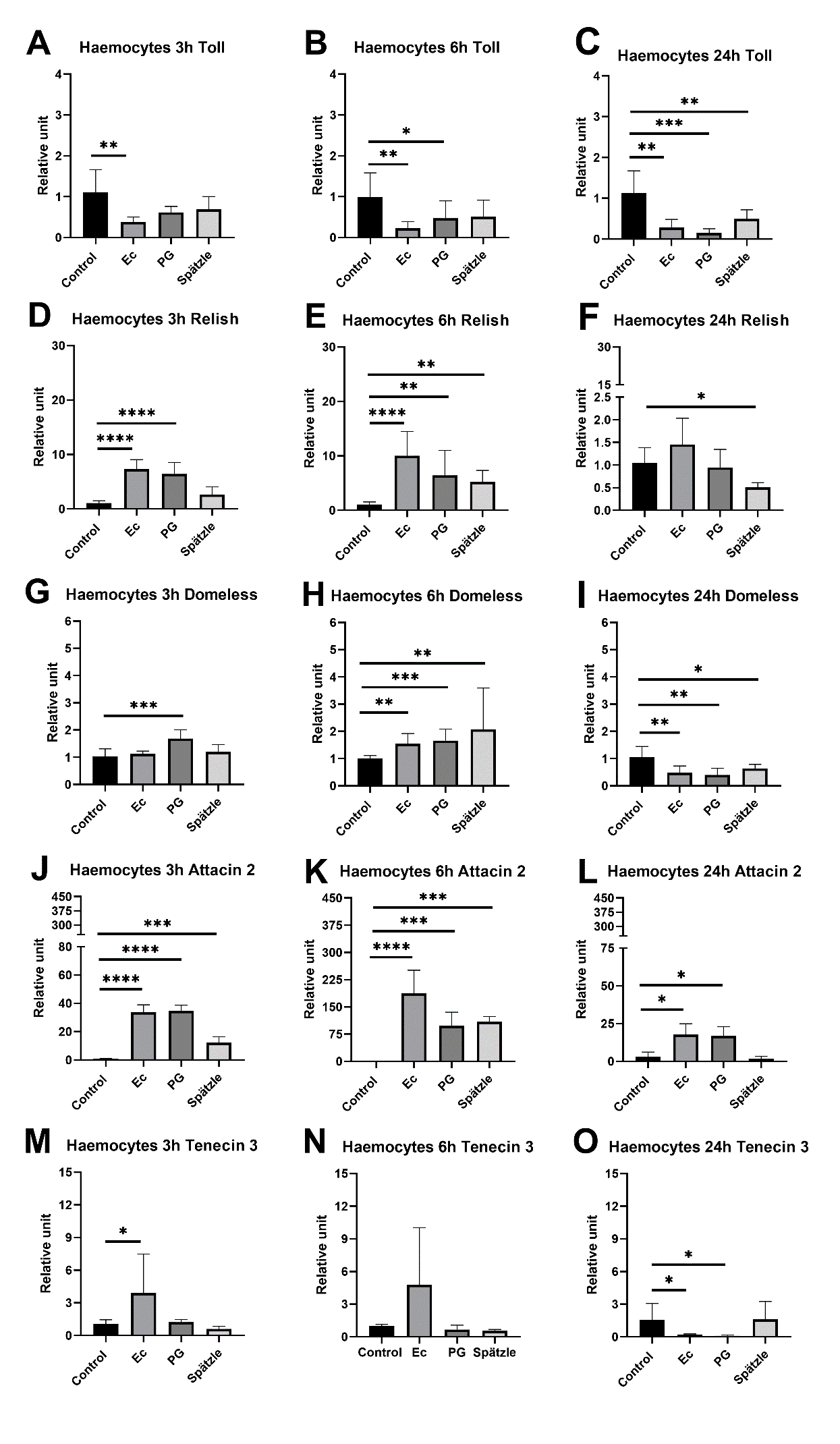

**Fig. S4.** Changes in the expression levels of immune-related genes (Toll (A-C), Relish (D-F), Domeless (G-I), Attacin 2 (J-L), Tenecin 3 (M-O)) in the haemocytes of *T. molitor* after activation of the immune system. The immune response was elicited by injection of *Escherichia coli* K16 (Ec), peptidoglycan from *Staphylococcus aureus* (PG), or Spätzle-like protein. Control – individuals injected with physiological saline. Due to the dynamic nature of the immune response, samples were collected 3, 6 and 24 hours after immune system activation. The values are the means ± SDs. Asterisks indicate statistically significant changes compared to those in control individuals; **p*≤0.05, ***p*≤0.01, ****p*≤0.001, **** *p*≤0.0001.

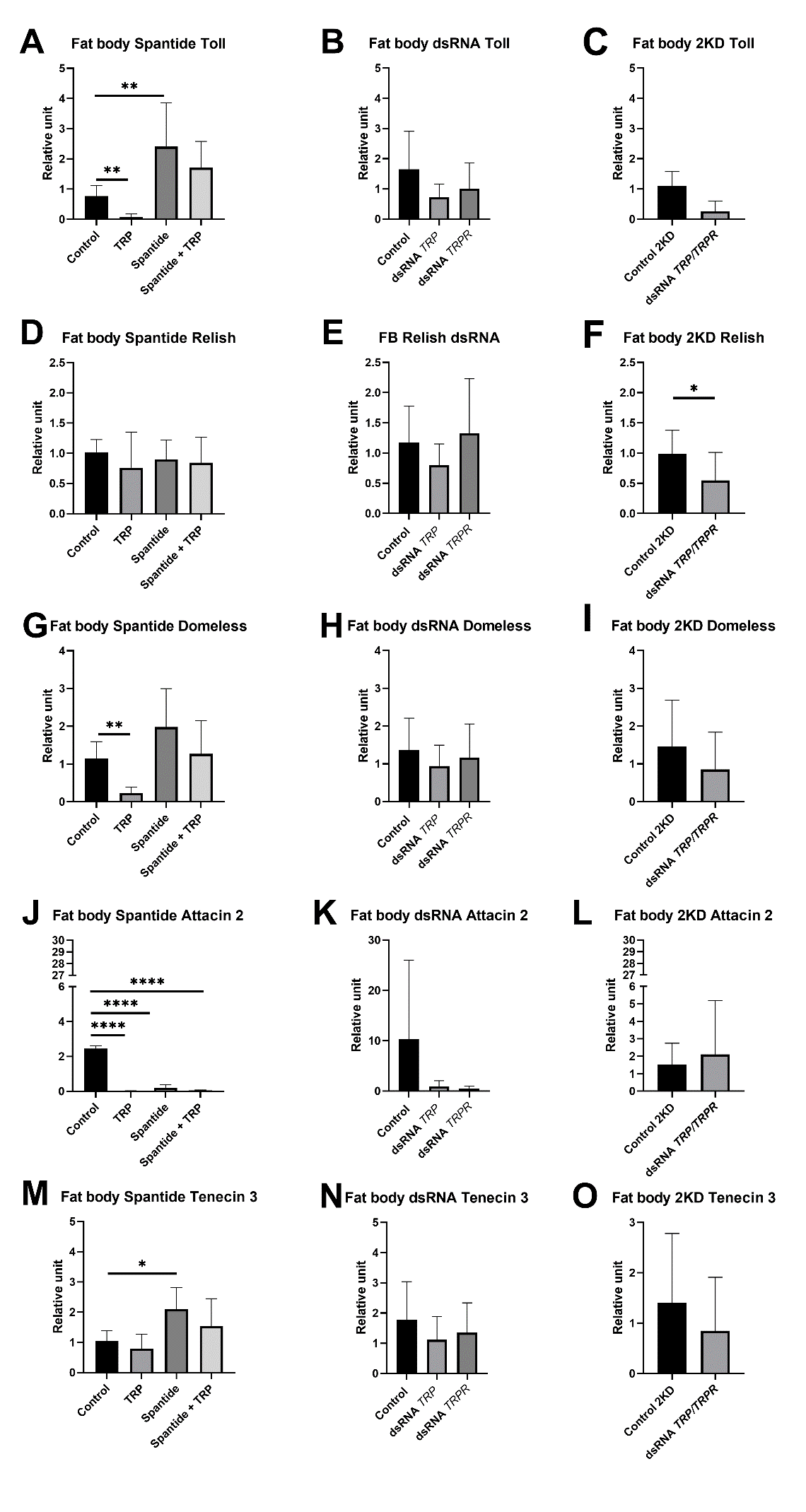

**Fig. S5.** Changes in the expression levels of immune-related genes (Toll (A-C), Relish (D-F), Domeless (G-I), Attacin 2 (J-L), Tenecin 3 (M-O)) in the fat body of *T. molitor* after application of physiological saline (control), Tenmo-TRP-7 (TRP) at a concentration of 10^-5^ M, Spantide II at a concentration of 10^-3^ M, and a mixture of Tenmo-TRP-7 and Spantide II. Also after injection of dsRNA targeted genes encoding lysozyme in *Galleria mellonella* (*LysGm*, Control) or genes related to TRP signalling (genes for TRP precursor (dsRNA TRP) or receptor (dsRNA TRPR). In addition, the double-knockdown of TRP and TRPR was analyzed. Control for the double-knockdown experiment (Control 2KD) was the injection of the double dose of dsRNA-targeted *LysGm*. The values are the means ± SDs. Asterisks indicate statistically significant changes compared to those in control individuals; **p*≤0.05, ***p*≤0.01, ****p*≤0.001, **** *p*≤0.0001.

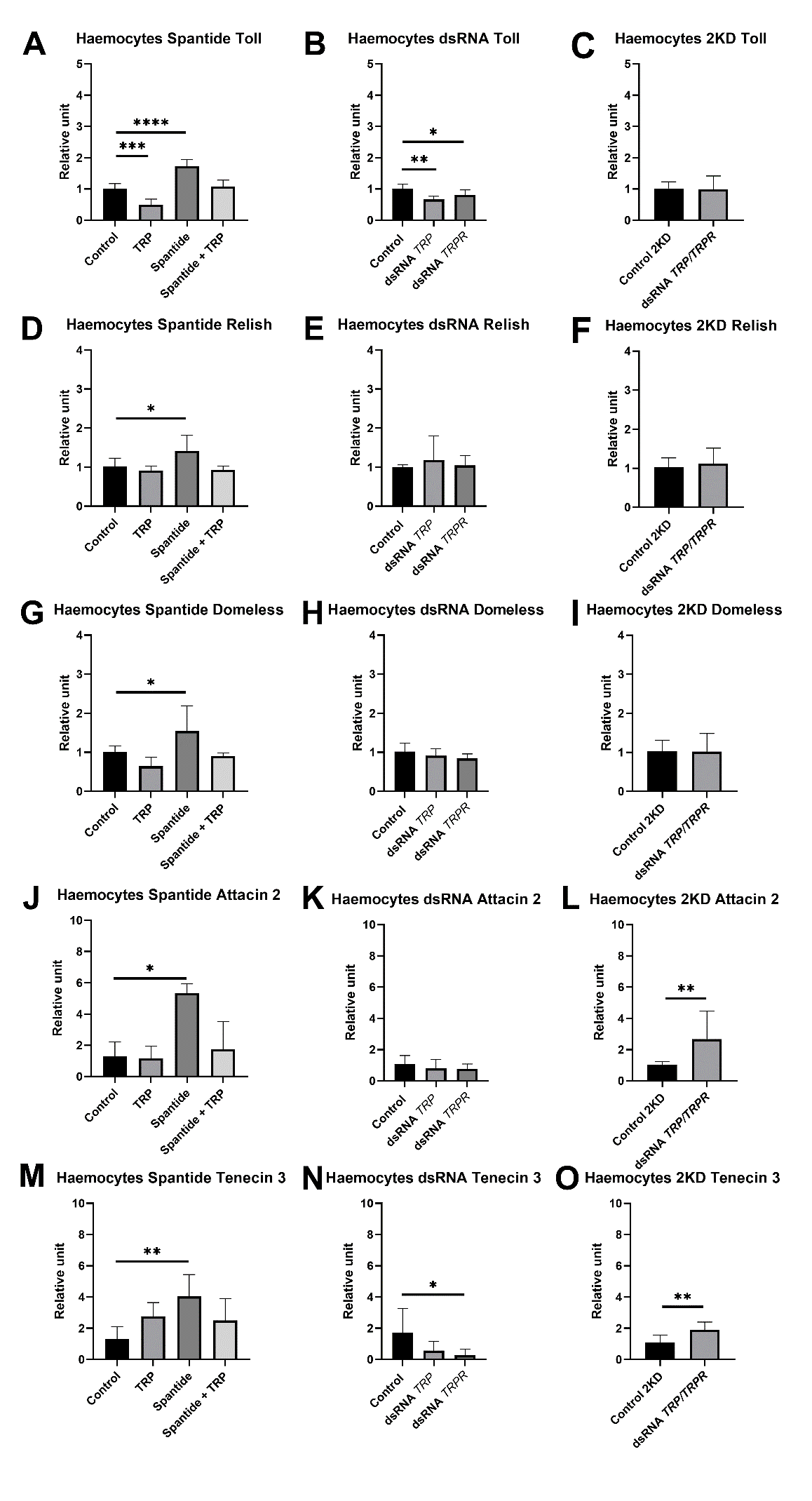

**Fig. S6.** Changes in the expression levels of immune-related genes (Toll (A-C), Relish (D-F), Domeless (G-I), Attacin 2 (J-L), Tenecin 3 (M-O)) in the haemocytes of *T. molitor* after application of physiological saline (control), Tenmo-TRP-7 (TRP) at a concentration of 10^-5^ M, Spantide II at a concentration of 10^-3^ M, and a mixture of Tenmo-TRP-7 and Spantide II. Also after injection of dsRNA targeted genes encoding lysozyme in *Galleria mellonella* (*LysGm*, Control) or genes related to TRP signalling (genes for TRP precursor (dsRNA TRP) or receptor (dsRNA TRPR). In addition, the double-knockdown of TRP and TRPR was analyzed. Control for the double-knockdown experiment (Control 2KD) was the injection of the double dose of dsRNA-targeted *LysGm*. The values are the means ± SDs. Asterisks indicate statistically significant changes compared to those in control individuals; **p*≤0.05, ***p*≤0.01, ****p*≤0.001, **** *p*≤0.0001.

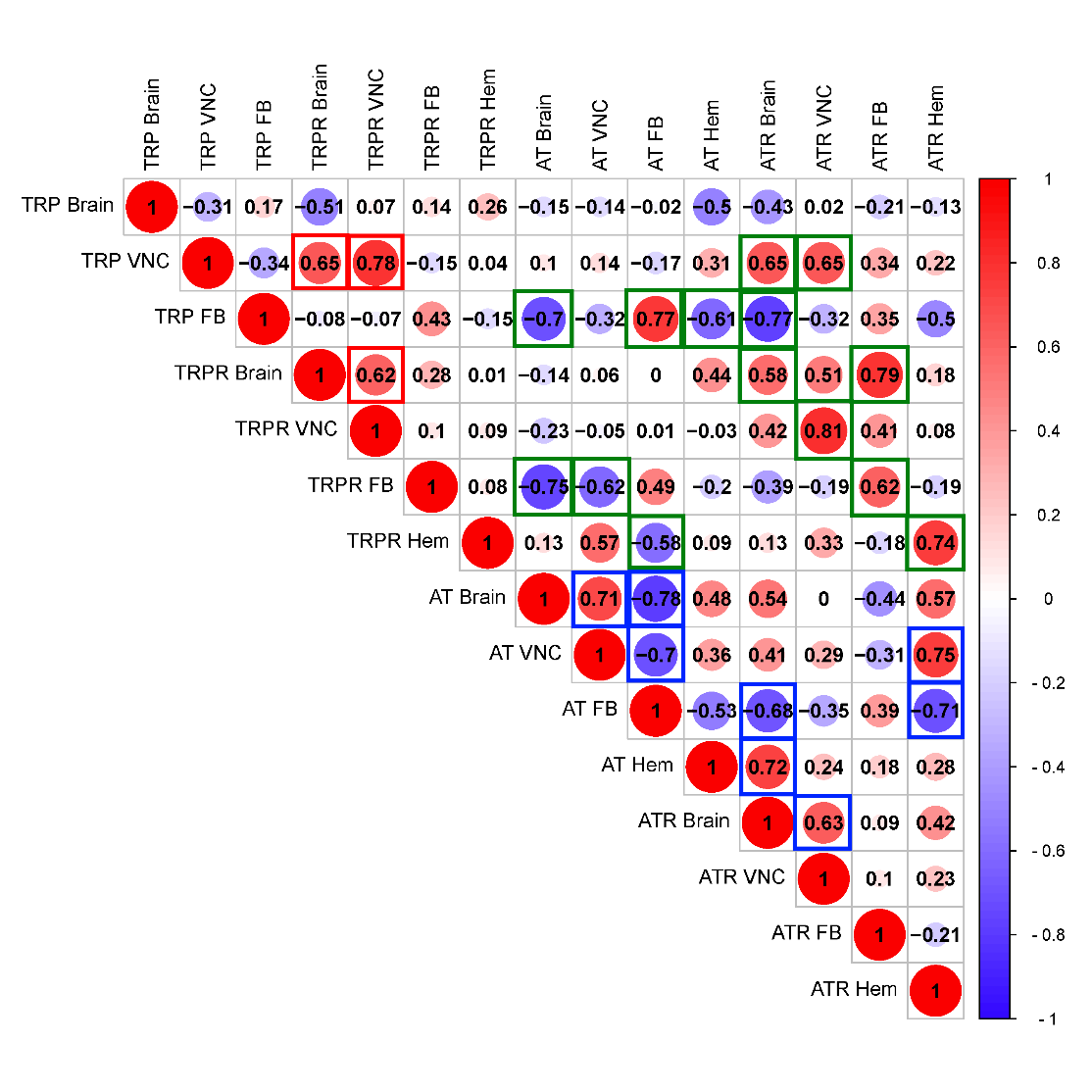

**Fig. S7.** Correlation of expression level of *TRP* and *TRPR* in different tissues/cells during activation of *Tenebrio* immune system with the expression level of genes encoding allatotropin precursor (AT) and receptor (ATR). Red squares – significant correlations associated with the TRP system presented in this article. Blue squares – significant correlations related to AT system (data previously presented by Konopińska et al. (2024)). Green squares – significant correlations between TRP and AT systems. VNC – ventral nerve cord; FB – fat body; Hem – haemocytes. To estimate the correlation of the data, the Pearson correlation coefficient method was used. The matrix was generated using SRplot software (https://www.bioinformatics.com.cn/srplot). Dot size – the level of the r value. The r value is presented in the middle of the dot. Different colors indicate different *r* values. Red shading indicates positive correlations (r > 0) and blue shading indicates negative correlations (r < 0). Red squares indicate statistically significant correlations (*p* ≤ 0.05).
